# Caught in Motion: Human NTHL1 Undergoes Interdomain Rearrangement Necessary for Catalysis

**DOI:** 10.1101/2021.08.13.456075

**Authors:** Brittany L. Carroll, Karl E. Zahn, John P. Hanley, Susan S. Wallace, Julie A. Dragon, Sylvie Doublié

## Abstract

Base excision repair (BER) is the main pathway protecting cells from the continuous damage to DNA inflicted by reactive oxygen species. BER is initiated by DNA glycosylases, each of which repairs a particular class of base damage. NTHL1, a bifunctional DNA glycosylase, possesses both glycolytic and ß-lytic activities with a preference for oxidized pyrimidine substrates. Defects in human NTLH1 drive a class of polyposis colorectal cancer. We report the first X-ray crystal structure of hNTHL1, revealing an open conformation not previously observed in the bacterial orthologs. In this conformation, the six-helical barrel domain comprising the helix-hairpin-helix (HhH) DNA binding motif is tipped away from the iron sulphur cluster-containing domain, requiring a conformational change to assemble a catalytic site upon DNA binding. We found that the flexibility of hNTHL1 and its ability to adopt an open configuration can be attributed to an interdomain linker. Swapping the human linker sequence for that of *Escherichia coli* yielded a protein chimera that crystallized in a closed conformation and had a lower binding affinity for lesion-containing DNA. This large scale interdomain rearrangement during catalysis is unprecedented for a HhH superfamily DNA glycosylase and provides important insight into the molecular mechanism of hNTHL1.

## INTRODUCTION

The base excision repair (BER) pathway recognizes, and repairs oxidized or alkylated DNA lesions, as well as mispaired uracil or thymine bases (1–3). This repair pathway entails a “hand off” of the DNA substrate in sequential steps (4,5). DNA glycosylases initiate the BER pathway by probing, recognizing, and excising DNA lesions. These enzymes are classified into two groups, monofunctional and bifunctional. The monofunctional enzymes possess only glycosylase activity, leaving an abasic site upon removal of the DNA lesion, whereas the bifunctional glycosylases have an additional lyase activity, which nicks the DNA backbone either once β-elimination, leaving a 3’-aldehyde; as is the case for endonuclease III (Endo III, Nth), or twice (β,δ-elimination, leaving a 3’-phosphate), thus creating a single-strand break (SSB) (6,7). Apurinic endonuclease 1 (APE1) or polynucleotide kinase (PNK) process the respective products, leaving a free 3’ hydroxyl for DNA polymerase β (Pol β) to insert the correct nucleotides using the undamaged strand as template. The nick is subsequently sealed by the DNA ligase IIIα (LIGIII)-X-ray repair cross complimenting 1 (XRCC1) complex (8). DNA glycosylases have a high affinity for their product and remain bound to the highly reactive abasic (AP) site until the hand-off to APE1 (9–13). The BER machinery likely does not scan the DNA as a complex but rather the transient interactions aid in recruitment of the downstream enzymes protecting the cells from potentially lethal intermediates (14,15).

Nth is a bifunctional DNA glycosylase with a marked preference for oxidized pyrimidine substrates (16). While both *Escherichia coli* Nth (EcoNth) and human NTHL1 (hNTHL1) glycosylases display a preference for excising lesions opposite G, the human enzyme is slower than the bacterial enzyme on most substrates (17–19). The first bacterial Nth crystal structure, EcoNth, revealed that the enzyme consists of two globular α-helical domains, a six-helical bundle domain and a [4Fe4S] cluster domain. This crystal structure identified the first HhH motif-containing glycosylase and the first [4Fe4S] cluster in a DNA-binding protein (20). Later, the *Geobacillus stearothermophilus* Nth was crystallized in the presence of DNA in two forms, bound to the non-hydrolysable AP site analogue, tetrahydrofuran (THF), and as a covalently trapped intermediate. These structures showed that DNA binds along the cleft between the two domains, with the nucleophilic lysine located in the six-helical barrel domain and the aspartate that deprotonates the lysine located in the [4Fe4S] cluster domain. The Nth DNA-bound models also revealed extensive contacts of the [4Fe4S] domain to the DNA backbone (21).

The BER pathway has been implicated in the progression of cancers because mutations in BER enzymes can reduce the effectiveness of the variants to repair damaged DNA, resulting in mutations (22). Loss of *NTHL1* is a driver of adenomas and other tumour types (see review (23)). Whole exome sequencing has directly linked *NTHL1* to familial inherited colorectal cancer (CRC), and adenomatous polyposis (24,25). Multiple studies have found that *NTHL1*-associated polyposis (NAP) is associated with biallelic germline nonsense mutations that renders the DNA glycosylase inactive. Biallelic mutations have been found in 14 different tumour types, notably colorectal, breast and endometrial cancers (24,26–31). Weren *et al*. estimate the prevalence of NAP at 1 in 114,770 in individuals of European decent (32). More recently, a paper reported a 1.9% prevalence of *NTHL1* biallelic mutations in polyposis patients, which typically present as adenomas potentially with CRC, serrated polyps, and multi-tumour phenotypes (33). The NAP tumours revealed a strong C>T transition pattern (24,34). *NTHL1* deficiency was also identified as the root of COSMIC mutation signature 30, using human intestinal organoids (35). Signature 30 was also identified in some breast cancers and, retrospectively, the breast tumour in which signature 30 was identified was determined to be *NTHL1*-deficient (36). Recently, four more breast tumours, where signature 30 accounts for 80% of the mutations, had an *NTHL1* deficiency, suggesting that lack of *NTHL1* had driven the formation of these tumours (28). Transcriptome sequencing of a pancreatic neuroendocrine tumour revealed signature 30 and *NTHL1* loss, implicating *NTHL1* deficiency as a driver of another tumour type (37). It was also reported that overexpression of NTHL1 causes genomic instability (38).

The bacterial and eukaryotic homologs share limited sequence homology, around 30%, and notable differences in activity. Mammalian NTHL1 DNA glycosylases, including hNTHL1, harbour a disordered N-terminal extension, which encompasses nuclear and mitochondrial localization sequences; it is also a site of post-translational modifications and has been posited to play a part in protein-protein interactions (39–41). Additionally, hNTHL1 exhibits apparent positive cooperativity that has not been observed in the bacterial homologs (39,41). These differences, and the need for structural mapping of cancer variants, highlight the need for a crystal structure of the human enzyme. Here we report the first crystal structure of hNTHL1, which was captured in a novel open conformation. A large-scale conformational change would be necessary to accomplish catalysis in DNA. We show that a linker between the two domains is necessary for the open conformation and that the freedom of movement between the two domains facilitates the excision of thymine glycol (Tg) from DNA oligomers.

## MATERIAL AND METHODS

### Expression of hNTHL1

The protein constructs, hNTHL1Δ63 and the hNTHL1Δ63 chimera, were overexpressed from the pET30a vector in *E. coli* Rosetta2 DE3 pLysS cells (Novagen), using autoinduction as previously described (42). Briefly, cells were grown in Terrific Broth media supplemented with 5052 sugar mix, kanamycin, and chloramphenicol at 20°C for 60 hours. The cells were lysed by sonication at 4°C in 500mM NaCl, 20 mM Tris pH 8, 20 mM imidazole, 10% (v/v) glycerol, 3 mM β-mercaptoethanol, 1 mM PMSF. The cell lysate was cleared at 23,000 x g for one hour and then passed over a Ni-NTA resin using gravity flow. The protein was eluted using 5 column volumes (CV) of elution buffer, 100 mM NaCl, 20 mM Tris-HCl pH 8, 250 mM imidazole, 10% glycerol, and 3 mM β-mercaptoethanol. The protein was further purified over a heparin column (Cytiva), in 100 mM NaCl, 20 mM Tris-HCl pH 8, 20 mM imidazole, 10% glycerol, 1 mM TCEP with a 20 CV salt gradient (0.1 – 1M NaCl). The protein typically elutes around 300 mM NaCl. A Superdex 75 gel filtration column (Cytiva) was used for the final purification step with a buffer composed of 100 mM NaCl, 20 mM HEPES pH 8, 10% glycerol, and 1 mM TCEP. Proteins were concentrated to ~8 mg/ml using a 30,000 Dalton cut-off centrifugal filter unit (Amicon), flash frozen in LN_2_, and stored at −80°C.

### Selenomethionyl-protein purification

The pET30a hNTHL1Δ63 construct was overexpressed in *E. coli* Rosetta2 DE3 pLysS cells (Novagen), in minimal medium containing selenomethionine at 125 μg/mL, as described in (43). Briefly, the cells were grown to an optical density of 0.6 at 37°C, and then induced with 500 mM IPTG at 25°C for 4 hours. The protein was purified using the procedure described above.

### Purification of DNA substrates

The following DNA substrates were chemically synthesized (IDT): 35mer oligonucleotides were used: damaged strand 5’ TGTCAATAGCAAG(X)GGAGAAGTCAATCGTGAGTCT 3’ and complementary strand 5’ AGACTCACGATTGACTTCTCC(G/A)CTTGCTATTGACA 3’, where X is the DNA lesion, either tetrahydrofuran (THF) or thymine glycol (Tg). (G/A) represents the opposite base. Oligonucleotides were resuspended in 200 μl TE buffer and 800 μl formamide and purified by electrophoresis on a 30% polyacrylamide-urea gel (National Diagnostics), run at 55 Watts for 6 hours. The bands were cut out of the gel, crushed, and soaked overnight in 50 mM NaCl and 50 mM Tris-HCl pH 8. The gel pieces were filtered out and the solution was run over a SepPak C18 column (Waters). The oligonucleotides were eluted in 75% acetonitrile. The acetonitrile was evaporated using a Speedvac, and the oligonucleotides were resuspended in annealing buffer (50 mM NaCl, 50 mM Tris-HCl pH 8). The oligonucleotides were annealed at an equimolar ratio using a hot water bath at 95°C and allowed to cool to room temperature slowly in the water.

### Crystallization of hNTHL1

Initial hits for hNTHL1Δ63 were obtained in the Index screen HT (Hampton Research) condition F12 (0.2 M NaCl, 0.1 M HEPES pH 7.5, 25% PEG 3350) set in a 96-well sitting drop plate by the NT8 drop setter (Formulatrix). Both hNTHL1Δ63 and hNTHL1 Δ63 SeMet crystals were grown in 24-well hanging drop trays with a final protein concentration of 3.5 mg/ml, incubated at 18°C, and streak seeded 18 hours later. The reservoir solution ranged from 0.5-2% Polyethylene glycol 5,000 monomethyl ether (PEG 5K MME, Hampton Research), 75 – 100 mM NaCl, and 100 mM tricine pH 8.5. Long needle-like crystals were cut to ~600 μM and soaked for 30 minutes in a 1:1 solution of mother liquor and a cryoprotecting solution of 10% (w/v) PEG 5K MME, 10 mM NaCl, 50% (w/v) glycerol, 50 mM Tricine pH 8.5. The hNTHL1Δ63 chimera crystals were grown using the hanging-drop method at a final concentration of 3 mg/ml, incubated at 18°C, and streak seeded 3 hours later. The reservoir solution ranged from 0.5 - 2% Polyethylene glycol 6,000 (PEG 6K, Hampton Research), 55 mM NaCl, and 100 mM Tricine pH 8.3. Needle-like crystals grew to approximately 400 x 70 x 70 μm^3^ over 5 days. Crystals were soaked for 30 minutes in a 1:1 solution of mother liquor and a cryoprotecting solution composed of 10% (w/v) PEG 6K, 10 mM NaCl, 50% (w/v) glycerol, 50 mM Tricine pH 8.3.

### Data collection and processing of crystals

hNTHL1Δ63 crystals diffracted to 2.5 Å at the APS synchrotron (beamline 23-ID-B). Data were collected using the 20-micron beam tuned to 12 kEV. A complete dataset was obtained by using the vector function along the length of the crystal. The hexagonal (P63) crystals were integrated, scaled, and truncated using iMOSFLM, AIMLESS/POINTLESS, and CTRUNCATE (44–48). Data were collected on hNTHL1Δ63 SeMet crystals at the APS synchrotron (beamline 23-ID-B) at a peak wavelength of 0.9795Å. The crystals diffracted past 3.2 Å. Data were collected using a 20-micron beam and the vector function described above. The SeMet crystals were hexagonal (P63) and were processed using HKL2000. hNTHL1Δ63 chimera crystals diffracted to 2.1 Å at the APS synchrotron (beamline 23-ID-D) at 12 kEV. The vector function was again employed in order to obtain a complete dataset. The hNTHL1Δ63 chimera crystallized in the orthorhombic space group, P2_1_2_1_2_1_. Data were integrated, scaled, and truncated using iMOSFLM, AIMLESS/POINTLESS, and CTRUNCATE (44–48).

### Structure Solution and Refinement

hNTHL1Δ63 crystals were experimentally phased by Single-wavelength Anomalous Dispersion (SAD) methods. The selenium sites were identified by PHENIX autoSOL (49,50). The hNTHL1Δ63 model was refined using PHENIX and Translation/Libration/Screw (TLS) model (49,51). There is one molecule per asymmetric unit (ASU) with a Matthew’s coefficient of 3.32 and a solvent content of 63 %. The hNTHL1Δ63 model was refined using PHENIX (49,51–54) to a Rwork/Rfree of 19.96%/24.45% at 2.5 Å with 96.96% preferred and 3.04% allowed residues in the Ramachandran plot. The average B-factor is 88.6 Å^2^; the high B-factor is most likely due to the high solvent content and flexibility of the two domains. The following residues were built in the electron density map: 86-317. The hNTHL1Δ63 chimera dataset was solved by molecular replacement using hNTHL1Δ63 as a search model. The two domains were separated and searched for individually using PHENIX autoMR (49). There is one molecule per ASU, with a Matthew’s coefficient of 2.36 and 48 % solvent content. The hNTHL1Δ63 chimera model was refined using PHENIX (49,51–54) to a Rwork/Rfree of 18.12%/23.06% at 2.1 Å with 98.1% preferred, and 1.9% allowed residues in the Ramachandran plot. The average B-factor is 29.5 Å^2^. The following residues were built in the electron density map: 86-106, 111-304.

### Radiolabelling DNA

DNA oligonucleotides were radio-labelled with ^32^P at a ratio of 10% hot DNA to 90% cold DNA for the single-turnover experiments and 100% hot DNA for EMSAs. The damage-containing oligonucleotide was incubated with ^32^P γ-ATP and polynucleotide kinase for 30 minutes. The reaction was quenched with 25 mM EDTA, and heat inactivated for 1 min at 95°C. The DNA was cleaned up by either ethanol precipitation for the 10% hot oligonucleotide, or a G-50 Probequant desalting column (Cytiva) for the 100% hot labelling. The damaged oligo and complementary strand were brought up to a final concentration of 250 nM in a DNA annealing buffer (10 mM Tris-HCl pH 8 and 50 mM NaCl) and annealed in 1:1 ratio in a 95°C water bath and allowed to cool slowly to room temperature.

### Single-Turnover Kinetics Experiments

To perform kinetics experiments under single-turnover conditions 100 nM of hNTHL1Δ63 or hNTHL1Δ63 chimera was added to a solution of 20 nM of radiolabelled DNA containing Tg:A, 2 mg/ml BSA (New England Biolabs), 10 mM Tris-HCl pH 8, 75 mM NaCl, and 1 mM DTT at 37°C. Time points were taken at 0, 0.25, 0.5, 1, 2, 3, 4, 5, 8, 10, 20, 30, 45 minutes and quenched in either equal volume of formamide stopping dye (95% formamide, bromophenol blue, xylene) or 0.1 M NaOH, and boiled for 5 min before adding equal volume formamide stopping dye. Samples were loaded onto a 12% sequencing gel and run at 55 watts for 1 hour. The gels were dried and exposed on phosphorescence screens (Kodak) and scanned using a STORM imager (GMI). Phosphorescence was quantified using Quantity One. The data were fit to the equation Y=Y0 + (Plateau-Y0)*(1-exp(-K*x)) using Graph Pad Prism (GraphPad Prism version 8.1.2 for Windows, GraphPad Software, La Jolla California USA).

### Electrophoretic Mobility Shift Assay

Electrophoretic Mobility Shift Assays (EMSAs) were performed with varying concentrations of hNTHL1Δ63 or hNTHL1Δ63 chimera from 0-1.75 nM, and 50 pM of radiolabelled THF:G DNA. The reactions were incubated in the same buffer listed above with 10% glycerol for 30 minutes at room temperature. The samples were run on a 6% non-denaturing polyacrylamide gel (AMERIBIO) at 4°C for 2 hours at 150 Watts. The gels were dried using a heater and vacuum pump and exposed overnight on phosphor screens. The gels were scanned using a STORM imager (GMI) and quantified with Quantity One (BioRad).

### Stopped-flow tryptophan fluorescence

Tryptophan fluorescence experiments were conducted on a stopped-flow SX-20 instrument (Applied Photophysics) with samples excited at 280 nm and emission filtered with a 350 nm long-pass filter at 37 °C. Data were collected using the pretrigger setting for 60 s. Artifacts from the initial flow and mixing of the solutions and instrument dead time were accounted for. Each trace reported is an average of multiple traces. The glycosylase reaction buffer was used: 10 mM Tris-HCl pH 8, 75 mM NaCl, 1 mM DTT. The final mixture contained 2 μM hNTHL1Δ63 or hNTHL1Δ63 chimera and 2 μM DNA substrate (same sequence as above).

### Selection of single nucleotide variants

We used the Genomic Data Commons (GDC) Application Programming Interface (API) to select subjects in The Cancer Genome Atlas (TCGA) that had at least one normal and one cancer sample. For the normal samples we used blood-derived normal, buccal cell normal, and solid tissue normal samples. Tumor samples were solid tumor samples. Of the 10,999 subjects in the TCGA, 9,962 met these requirements. After selecting the subjects, we used the GDC API to perform BAM slicing for *NTHL1*. Using VarScan 2.4.4 and the GRCh38 as a reference, we tested for germline mutation using every possible combination of tumor and normal samples for each subject and thresholds of minimum variant frequency > 0, p-value < 0.1 and a minimum depth of eight sequence reads covering each variant. Once the variants were called for each subject, we selected the exon chromosome positions that had mutations in at least 10 subjects. For those exons with at least 10 subjects, we determined if the mutation was a missense, nonsense, or silent and recorded the number of cancers associated with each mutation. The potential pathogenicity of the mutations was assessed using REVEL, where higher scores are correlated with a higher likelihood of being disease-causing (55).

## RESULTS

### Human hNTHL1 structure adopts an open conformation

We present the first crystal structure of hNTHL1, which was solved using single-wavelength anomalous dispersion (SAD) to 2.5 Å using the peak anomalous signal of selenium with a resulting R-work/R-free of 19.96%/24.45% (Table 1, Figure 1). For crystallization, we selected hNTHL1Δ63, a construct lacking the first 63 residues (previously described as hNTHL1Δ55 (39,40) due to a discrepancy in the identity of the initiation methionine) because the flexible N-terminal region appeared to hinder crystal formation of full-length hNTHL1. The crystal structure contains 227 residues from hNTHL1 (aa 86-312), in addition to the 6xHis-Tag, forming a total of 10 α helices. Like its bacterial homologs, hNTHL1 comprises two globular domains: a six-helical bundle domain, which contains a helix-hairpin-helix (HhH) DNA-binding motif and a helical domain containing a [4Fe4S] cluster domain (Figure 1)(20,21,56) (57). Both the N- and C-termini of hNTHL1 reside in the [4Fe-4S] cluster domain; the two domains are therefore connected by two linkers: linker 1 (aa 104-125) joins the [4Fe-4S] cluster domain to the six-helical bundle domain and linker 2 (aa 230-240) connects the six-helical bundle domain to the [4Fe4S] cluster domain (Figure 1). Surprisingly, our hNTHL1Δ63 model reveals a novel open conformation (Figure 2A), which is not observed in any of the bacterial Nth glycosylases previously crystallized without DNA: *Escherichia coli* Nth (EcoNth) (PDB ID 2ABK (58), RMSD 7.4 Å, calculated with PyMOL (59) and *Deinococcus radiodurans* Nth (DraNth) (PDB IDs 4UNF & 4UOB (57), RMSD 8.4 Å & 10.3 Å, respectively). The unliganded bacterial Nth structures have a domain orientation similar to that of the DNA-bound *Geobacillus stearothermophilus* Nth (GstNth) (PDB ID 1ORN (21), RMSD between GstNth and EcoNth is 1.9 Å).

**Figure 1:**
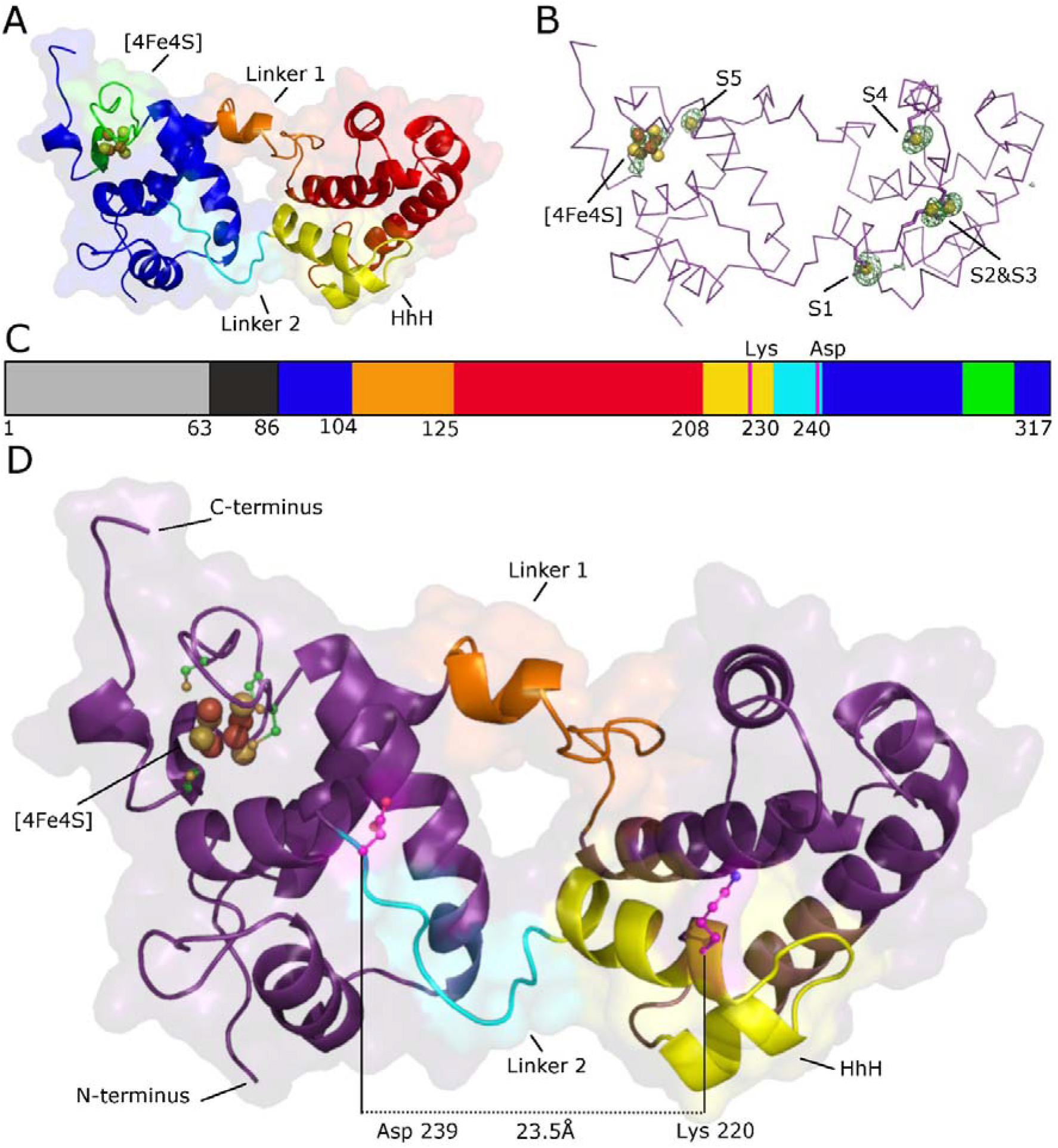
The active site of human NTHL1 is unassembled in the absence of DNA. A) An overview of hNTHL1Δ63 coloured by domain and motifs. The [4Fe4S] domain is shown in blue, the six-helical bundle domain in red. Linker 1 is coloured orange, the HhH DNA binding motif in yellow, linker 2 in cyan, and the [4Fe4S] cluster in green. B) The peaks (green mesh) from an anomalous difference Fourier map calculated with the peak selenium data set overlay nicely on the sulphurs (spheres) from the five methionine residues in the hNTHL1Δ63 model. There is a sixth anomalous peak, which corresponds to the [4Fe4S] cluster. C) A cartoon representation of the hNTHL1 domain organization, from N-to C-terminus, with the same colour code as in panel A: missing residues (light grey not included in the protein construct, dark grey included in the protein construct but not built in the model), six-helical bundle (red), linker 1 (orange), HhH (yellow), linker 2 (cyan), [4Fe4S] domain (blue); the catalytic residues, Lys 220 and Asp 239 (magenta), and the [4Fe4S] cluster (green). D) A detailed view of hNTHL1 Δ63 highlighting the unassembled active site illustrates the distance between the catalytic Lys and Asp (23.5 Å). Same colour code as in C.

**Table 1:** hNTHL1 Data Processing and Refinement Statistics. Data collection and refinement statistics for the hNTHL1Δ63, hNTHL1Δ63 chimera, and hNTHL1Δ63 SeMet crystals. The highest resolution shell is shown in parentheses.

TABLE AND FIGURE LEGENDS
|  | hNTHL1Δ63 | hNTHL1Δ63 chimera | hNTHL1Δ63 SeMet |
| --- | --- | --- | --- |
| **PDB ID** | 7RDS | 7RDT |  |
| **Data Collection** |  |  |  |
| Beamline | APS 23-ID-B | APS 23-ID-D | APS 23-ID-B |
| Space group | P 6<sub>3</sub> | P 2<sub>1</sub> 2<sub>1</sub> 2<sub>1</sub> | P 6<sub>3</sub> |
| Cell dimensions |  |  |  |
| a,b,c (Å) | 124.8 124.8 42.25 | 42.37 71.68 86.52 | 125.0, 125.0, 42.4 |
| α,β,γ (°) | 90 90 120 | 90 90 90 | 90 90 120 |
| Resolution range (Å) | 40.85- 2.5 | 38.05 - 2.1 | 40 - 3.2 |
| R-pim (%) | 6.8 (62) | 5.3 (21) | 10.6 (55.6) |
| CC1/2 | 99.5 (57.3) | 99.5 (84.3) | 97.6 (16.7) |
| I/sigma | 8.3(1.8) | 8.9 (3.4) | 12.1 (2.87) |
| Completeness (%) | 100 (100) | 99.7 (99.3) | 100 (100) |
| Multiplicity | 22.2 (22.3) | 4.1 (4.0) | 8.4 (8.5) |
| Wavelength (Å) | 1.033 | 1.033 | 0.9795 |
| Wilson-B factor (Å<sup>2</sup>) | 50.72 | 22.48 |  |
| **Refinement** |  |  |  |
| Resolution range (Å) | 40.85 - 2.5 | 38.05- 2.1 |  |
| R-work/R-free (%) | 19.96/24.45 | 18.12/23.06 |  |
| No. reflections | 25,423 | 28,900 |  |
| No. atoms |  |  |  |
| Protein | 1,756 | 1,725 |  |
| Ligands | 8 | 8 |  |
| Water | 115 | 200 |  |
| RMS Deviations |  |  |  |
| Bond lengths (Å) | 0.002 | 0.002 |  |
| Bond angles (°) | 0.431 | 0.471 |  |
| B-factor (Å<sup>2</sup>) | 88.64 | 29.49 |  |
| Protein | 89.34 | 28.83 |  |
| Ligands | 73.19 | 28.03 |  |
| Water | 78.77 | 35.11 |  |
| Ramachandran |  |  |  |
| Preferred (%) | 96.96 | 98.10 |  |
| Allowed (%) | 3.04 | 1.90 |  |
| Outlier (%) | 0 | 0 |  |
| Rotamer outlier | 1.06 | 0.56 |  |
| Clashscore | 6.64 | 2.05 |  |
| Number of TLS groups | 3 |  |  |

### The flexible linker is necessary for the open state

In the open state, the strictly conserved catalytic residues, Lys 220 and Asp 239, which are located ~5 Å apart in the bacterial enzymes, are found ~23 Å apart (Figure 1). Therefore, hNTHL1 must undergo a conformational change in order for catalysis to occur (17,19,60–62). This finding was unexpected as the bacterial homologs were captured in a similar closed conformation, in the presence and absence of DNA (Figure 1). We noted that *E. coli* Endonuclease 8 (EcoNei), a DNA glycosylase from the Fpg/Nei family, undergoes a conformational change between the unliganded and DNA-bound forms (Figure S1). EcoNei, like hNTHL1, also harbours a flexible hinge region (63). Based upon this precedent, we investigated the role of linkers in hNTHL1. A multiple sequence alignment with mammalian, lower eukaryotes, and prokaryotes created using MUSCLE showed that linker 1 is not highly conserved (64). To visualize the conservation of Nth residues, we used CONSURF (65) to generate a multiple sequence alignment of 150 orthologs, and colour the residues by conservation level from red to blue (conserved to variable). As mentioned above, linker 1 exhibits a degree of divergence, with a single conserved turn, whereas linker 2 is more conserved across species (Figure 2). A simplified sequence alignment shows an eleven-amino acid insertion compared to bacterial Nth sequences in the linker 1 region: seven residues extend the loop to 21 residues and four residues extend helix B in hNTHL1. In contrast, 14 residues compose the linker in *E. coli* (Figure 2, orange). Linker 2 is more conserved, with a single residue insertion in the eukaryotic NTHL1 (Figure 2). To test whether the open conformation in hNTHL1 was due to the increased flexibility imparted by the extended linker 1, we engineered a chimera of hNTHL1Δ63 and EcoNth by replacing 16 residues (residues 110-125) of the human linker with the shorter EcoNth linker (residues 21-28) (hNTHL1Δ63 chimera, Table 2). Since the length of linker 2 was quite similar across bacterial and mammalian sequences, we hypothesized that the increased flexibility in the human enzyme was due to linker 1, and linker 2 was therefore not altered in this report.

**Figure 2:**
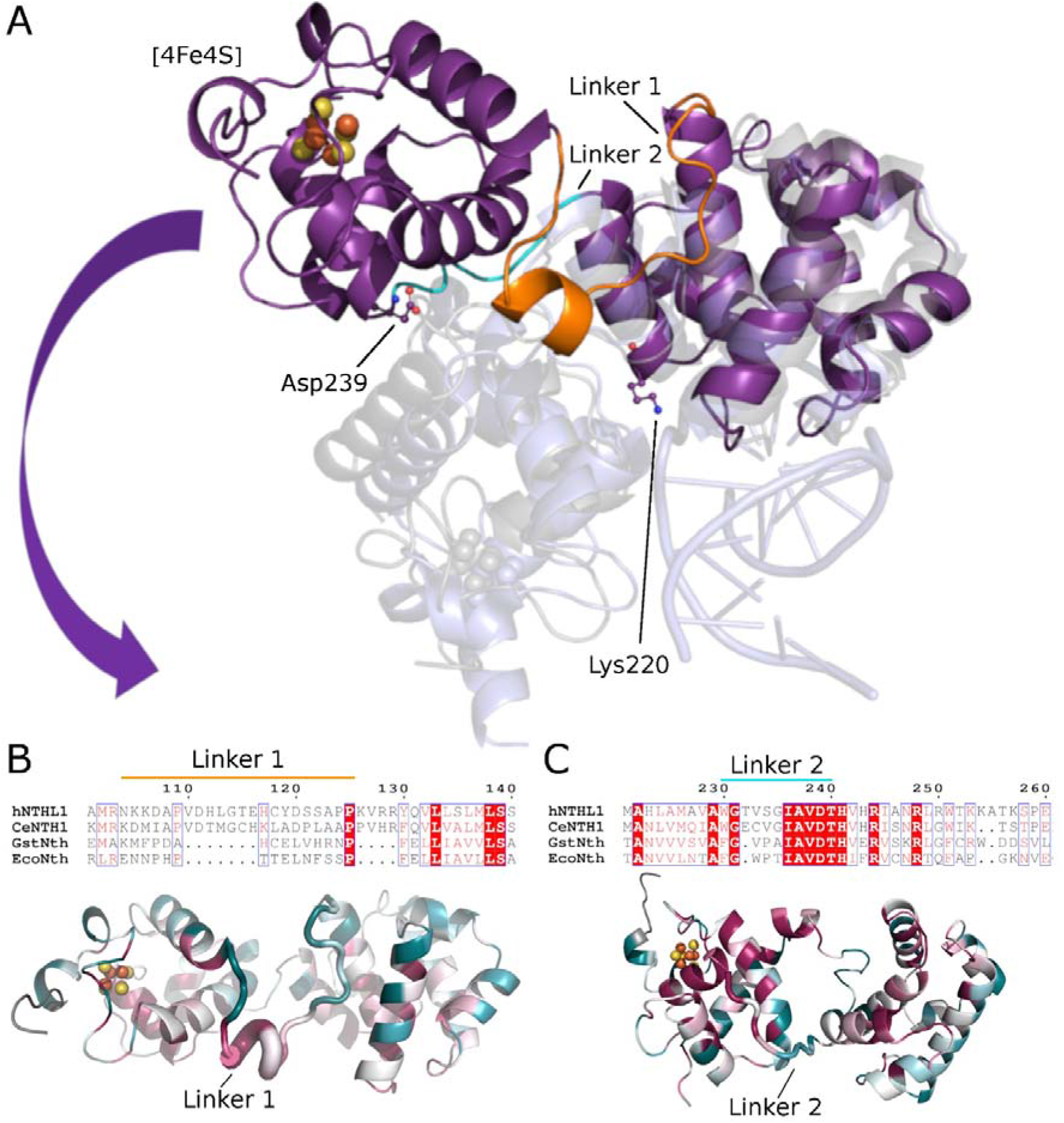
An extended linker 1 in hNTHL1 creates a flexible hinge region. A) The unexpected domain orientation of hNTHL1 Δ63 (purple) relative to *G. stearothermophilus* (PDB ID code 1ORN, transparent blue (21)) became evident from a superposition of the two orthologs. Linker 1 is shown in orange and linker 2 is shown in cyan. B) A multiple sequence alignment (64,80) between human, *C. elegans, E. coli*, and *G. stearothermophilus* confirms the variability of linker 1 (orange line), where eukaryotic NTHL1 has a seven amino acid insertion in addition to four additional residues extending the helix immediately after linker 1. Linker 2 is more conserved, with a single residue insertion in the eukaryotes. C) The CONSURF (65) representation of sequence homology (red, conserved and blue, variable) shows a higher degree of variability in linker 1 than in linker 2.

**Table 2:** Nth linker 1 sequences. The linker sequences for hNTHL1, EcoNth, and hNTHL1Δ63 chimera are listed. The human linker (linker 1) was substituted with the corresponding EcoNth linker. The last four residues (KVRR) are an extension of helix B and were therefore left in the hNTHL1Δ63 chimera to not disrupt local secondary structure elements.

| Construct | Linker Sequence |
| --- | --- |
| **hNTHL1Δ63** | VDHLGTEHCYDSSAPKVRR |
| **hNTHL1Δ63 chimera** | TTELNFSSPKVRR |
| **EcoNth** | TTELNFSSP |

We solved the crystal structure of hNTHL1Δ63 chimera to 2.1Å (R-work/R-free of 18.12%/23.06%) by molecular replacement using hNTHL1Δ63 as the search model (49,66). To determine the phases via molecular replacement, the six-helical bundle domain was placed prior to the [4Fe4S] domain. By searching with each domain individually, the domains were allowed the freedom necessary to rearrange relative to each other. The hNTHL1Δ63 chimera crystallized in a closed conformation, similar to unliganded EcoNth (RMSD 2.21 Å), DraNth (RMSD 2.77 Å & 1.84 Å), and GstNth bound to DNA (RMSD 1.40 Å). To examine the degree of rotation, we superimposed the open and closed models in COOT (67) using the six-helical bundle domain as a reference, and then measured the angle of rotation between three residues, Asp 239 hNTHL1Δ63, Lys 220 hNTHL1Δ63/hNTHL1Δ63 chimera, and Asp 239 NTHL1Δ63 chimera, calculating a ~85° rotation (Figure 3 A, S2). When comparing the WT to the chimera construct, the analogous catalytic aspartates were 23.6 Å apart, and the [4Fe4S] clusters were 38.7Å apart (Figure S2). We were unable to completely trace the hNTHL1Δ63 chimera linker itself because of disorder in the electron density map. Nevertheless, the shortened linker decreased the interdomain flexibility within the enzyme. DNA interacting regions are highly conserved while the outer edges of the enzyme display more variability, as visualized with CONSURF (65)(Figure S3). The closed conformation may be stabilized by interactions between the globular domains and linker 2: Arg 272 of the [4Fe4S] domain interacts with the backbone carbonyl of Val 233 of linker 2; a second hydrogen bond is formed between His 223 (six-helical bundle domain) and Ser 234 (linker 2); and the third interaction occurs between Glu 267 ([4Fe4S] domain) and the backbone amide of Leucine 236 (linker 2) (Figure 3). The refined B-factors, or temperature factors, provide a measure of the relative movement of atoms, where higher numbers (shown in warmer colours in the protein cartoon) signify increased movement (Figure 3). The reduced movement of the chimeric enzyme is reflected in the B-factors: not only is the average B-factor lower overall, but the [4Fe-4S] cluster domain has greatly reduced B-factors in the chimera construct relative to the open hNTHL1Δ63 structure.

**Figure 3:**
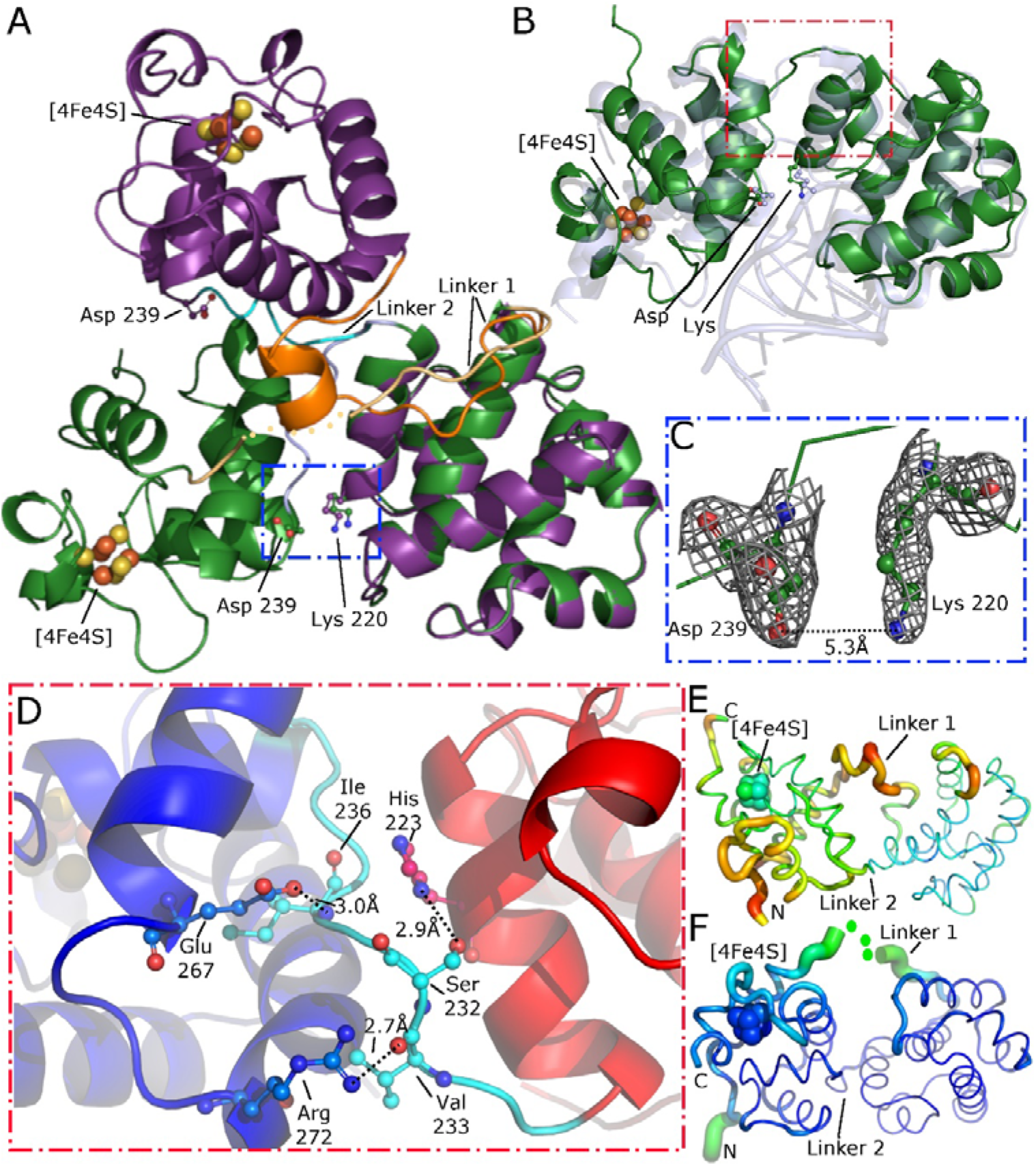
Shortening linker 1 in hNTHL1 chimera assembles the active site. A) Shortening linker 1 in hNTHL1Δ63 chimera (green) decreased the interdomain flexibility and yielded a closed domain orientation compared to hNTHL1Δ63 (purple). Linker 1 is shown in orange, and light orange for hNTHL1Δ63 and hNTHL1Δ63 chimera, respectively. Linker 2 is shown as cyan and light blue for hNTHL1Δ63 and hNTHL1Δ63 chimera, respectively. The catalytic residues (Lys 220 and Asp 239) are shown as sticks. B) Comparison of hNTHL1Δ63 chimera (green) to DNA-bound *G. stearothermophilus* Nth (transparent blue, PDB 1ORN (21)) reveals a similar domain orientation. C) Zooming in on the active site (blue dashed box) reveals a complete active site with the catalytic lysine and aspartate 5.3 Å apart (2F_O_-F_C_ map is shown as a grey mesh). This corresponds to the distances observed in the bacterial homologs. D) The closed conformation may be stabilized by interactions between the globular domains and linker 2. hNTHL1Δ63 chimera is shown in cartoon format with the highlighted residues shown as sticks; the [4Fe4S] domain is shown in blue, linker 2 in cyan, and the six-helical bundle domain in red. Arg 272 interacts with the backbone carbonyl of Val 233, a second hydrogen bond is formed between His 223 and Ser 234, and the third interaction occurs between Glu 267 and the backbone amide of Leu 236. B-factor analysis suggests that the shortened linker 1 reduces movement of hNTHL1. E) A putty representation of hNTHL1Δ63 coloured by B-factor shows that the [4Fe4S] domain has more movement than the six-helical bundle domain (A high B-factor signifies a higher degree of movement, indicated by warmer colours and a thicker putty). F) The same B-factor representation of hNTHL1Δ63 chimera shows decreased movement within the entire protein but especially in the [4Fe4S] cluster relative to the six-helical bundle domain.

### The closed conformation of hNTHL1Δ63 chimera restores the active site

In the hNTHL1Δ63 chimera model, the catalytic Asp and Lys are 5.3 Å apart. These residues occupy the identical orientation in the closed structures of homologs. Closure of the enzyme therefore restores the glycosylase active site to an apparent active configuration (Figure 3). Comparing these new structures of hNTHL1 to prior models of homologs suggests that the conformation change is essential for catalysis. These conformational changes observed in the hNTHL1Δ63 crystal structures are reminiscent of the conformational changes reported for EcoNei, which also closes upon binding DNA (63,68). In the apo-EcoNei structure, the C-terminal domain requires a reported ~50° rotation to form the closed ligand complex (63)(Figure S1). Another glycosylase of the Fpg/Nei family, Nei-like 2 (NEIL2) is also predicted to undergo a similar conformational change between the unliganded and bound forms (69).

### The hNTHL1Δ63 chimera has reduced DNA binding and activity

To determine if the hNTHL1Δ63 chimera retains activity we performed glycosylase activity assays under single-turnover conditions and electrophoretic mobility shift assays (EMSAs). In the glycosylase assays, the 5’ end of the DNA strand containing the lesions was radiolabelled with ^32^P. hNTHL1 nicks the DNA backbone as the damaged base is removed, resulting in a single product band with increased electrophoretic mobility on a PAGE gel. When the assays are quenched with NaOH the glycosylase activity is measured, as NaOH will resolve the Schiff base and reduce the AP site. Alternately, quenching with a formamide dye yields a single product band representing both glycosylase and lyase activity, because hNTHL1 must catalyse the resolution of the Schiff base and β-elimination. Based upon the published catalytic scheme for hNTHL1, k_2_ is the rate of base excision, and k_3_ is the rate of β-lyase activity (Figure S4) (60). We found that the hNTHL1Δ63 chimera retains some glycosylase activity but is impaired compared to the parental variant (Table 3, Figure 4A). The hNTHL1Δ63 chimera is highly deficient when it must provide the lyase reaction (Table 3, Figure 4B). It was not possible to calculate the rate constant k_3_ for the Tg:A substrate for the chimera construct because the data did not fit a one-phase exponential association model. The EMSA experiments showed that the hNTHL1Δ63 chimera has an impaired ability to shift DNA containing the non-hydrolysable AP site analogue, THF, compared to hNTHLΔ63 (Figure 4C). This finding suggests that the flexibility imparted by the longer linker is crucial for binding DNA. Taken together, these results indicate that when hNTHL1Δ63 is trapped in the closed conformation, DNA binding is impaired. The deficiency in glycosylase and lyase activity likely follow from this DNA binding defect. The reduction in catalytic activity is likely due to the impaired DNA binding, because when the chimeric DNA glycosylase is provided with additional time, it can cleave all the DNA substrate in the assay (Figure 4).

**Figure 4:**
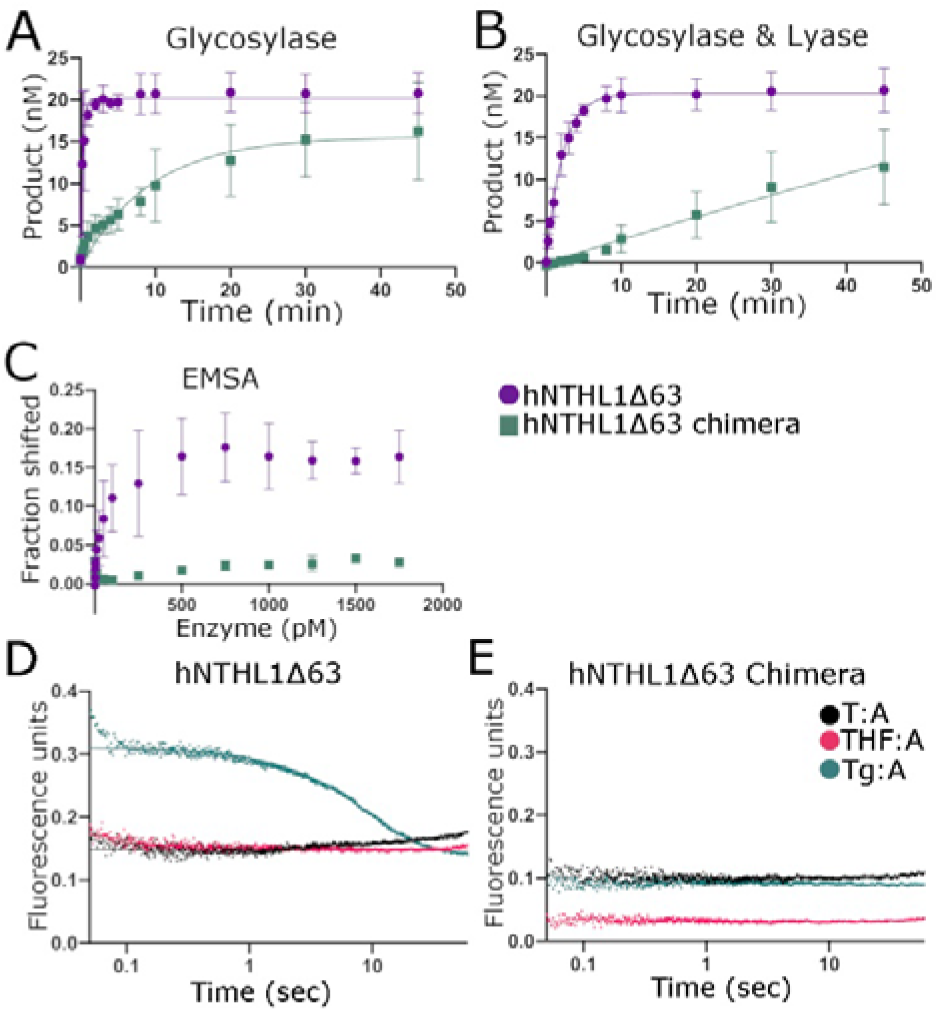
Interdomain flexibility is required for DNA binding. Single-turn over experiments using Tg:A and hNTHL1Δ63 and hNTHL1Δ63 chimera for (A) the β-elimination reaction and (B) only the glycosylase reaction. C) Quantification of the EMSA of hNTHL1Δ63, and hNTHL1Δ63 chimera with DNA containing THF:G shows that the chimera construct is unable to shift an appreciable amount of DNA. Stopped flow experiment measuring tryptophan fluorescence of hNTHL1Δ63 (D) and hNTHL1Δ63 chimera (E) incubated with T:A, THF:A, and Tg:A.

**Table 3:** Single-turnover experiments with hNTHL1Δ63 and hNTHL1Δ63 chimera. The rate constants for hNTHL1Δ63 and hNTHL1Δ63 chimera are reported in this table. The rate constant k_2_ is the observed rate of glycosylase only activity, quenched with NaOH. The rate constant k_3_ is the observed rate of β-elimination activity, quenched with formamide stopping dye. Experiments were repeated three times. The rate constants were calculated by fitting the data to a one-phase association exponential, Y=Y0 + (Plateau-Y0)*(1-exp(-K*x)). The hNTHL1Δ63 chimera has a significantly slower β-lytic activity on DHU:G and Tg:A, and glycosylase activity on Tg:A but not DHU:G.

|  | Tg:A |  |
| --- | --- | --- |
| | $k_2$ (min<sup>-1</sup>) | $k_3$ (min<sup>-1</sup>) |
| **hNTHL1Δ63** | 3.2 | 0.46 |
| **hNTHL1Δ63 Chimera** | 0.11 | n/a |

### The hNTHL1Δ63 chimera has reduced interdomain flexibility

To determine if a conformational change could be observed during DNA binding, we performed stopped-flow kinetics experiments and measured tryptophan emission at 490 Abs with hNTHL1Δ63 and hNTHL1Δ63 chimera, in the presence of Tg:A, THF:A, or undamaged DNA. Figure S5 shows the location of the seven tryptophan residues present in hNTHL1Δ63. We observed a conformational change for hNTHL1Δ63 when processing Tg:A, but not THF:A or undamaged DNA (Figure 4D). There was no conformational change detected with the hNTHL1Δ63 chimera on any of the DNA substrates (Figure 4D). The fact that we observed a change in the hNTHL1Δ63 conformation during lesion processing, but not in the hNTHL1Δ63 chimera, suggests that a conformational change occurs during lesion removal that is dependent upon the presence of a flexible linker. Moreover, the conformational change that was observed in hNTHL1Δ63 was seen only in the context of a lesion, Tg, suggesting that the structural changes resulting in a difference in tryptophan environment happen during or after base cleavage.

## DISCUSSION

The first crystal structure of human NTHL1 reveals an open conformation that had not been observed previously in any of the bacterial Nth homologs, or other members of the HhH family such as OGG1 or MutY. In the open conformation the DNA glycosylase is catalytically inactive as the two active site residues (K220 and D239) are much too far apart, 23.5 Å, to perform a nucleophilic attack. Thus, a conformational change must occur upon DNA binding to assemble the active site. We showed that the interdomain rearrangement requires an extended linker in the hNTHL1 enzyme. With the substitution of the *E. coli* linker in hNTHL1Δ63, hNTHL1Δ63 chimera crystallized in the closed form, like that of bacterial Nth, assembling the active site (16,20,21,56–58). Even though the active site is assembled, the hNTHL1Δ63 chimera exhibits a severely decreased glycosylase activity and reduced DNA binding affinity. This finding suggests that the movement between the two domains is critical for DNA binding *in vitro*.

Mapping residues onto a three-dimensional structure is crucial when designing and interpreting mutagenesis studies, and for understanding the potential outcome of cancer variants. A series of hNTHL1 point mutations can now be interpreted considering the new crystal structures. Robey-Bond *et al*. investigated several residues which were deemed important for catalysis in *E. coli* and yet had a different amino acid residue at the analogous position in mammalian NTHL1 (19), including Asn279, Gly280, and Gln287 (19). Our crystal structure of hNTHL1 shows that Asn 279 and Gly 280 are located near linker 2, with the former contacting the backbone carbonyl of Ala 137 in that linker. hNTHL1 Gln 287 is predicted to contact the DNA backbone. The analogous residue in GstNth, Arg186, contacts the DNA backbone at the site of the lesion, and hydrogen bonds Glu 24 to linker 1. Additionally, we were able to map germline hNTHL1 single-nucleotide variants (SNVs) from the TCGA dataset (T able S1), onto the open and closed hNTHL1 models. The curated SNVs were predicted to be deleterious using REVEL (55). Interestingly, these mutations appear to cluster near the two linker regions, and therefore could potentially affect the predicted conformational change (Figure S6). While none of the SNVs are close enough to contact the DNA, Ile 176 is part of the HhH motif and Thr 289 is close to the [4Fe4S] cluster.

DNA glycosylases were generally thought to be relatively rigid enzymes, with modest interdomain changes, as the apo- and DNA-bound forms are nearly identical in numerous X-ray crystal structures where both DNA-bound and unbound models are available (Figure S1). There are now several examples of glycosylases that show interdomain rearrangement when comparing the unliganded to the DNA-bound form. Human uracil DNA glycosylase (UDG) undergoes a 10° conformational change, in which the two globular domains move together and “pinch” the DNA, causing a 45° kink in the DNA backbone (70). EcoNei has a much more dramatic 50° interdomain rotation upon binding DNA (63). In the same H2TH family as EcoNei *Neisseria meningitidis* formamidopyrimidine DNA glycosylase (Fpg) was shown to undergo a 22° conformational change upon binding DNA (69). Additionally, the unliganded mammalian NEIL2 crystalized in an open conformation with a large conformational change upon binding DNA shown using small angle X-ray scattering(69). NTHL1, is the first reported member of the HhH family of DNA glycosylases to show structural evidence of a large conformational change and joins the UDG and H2TH glycosylase families in this respect (63,70). A structure based alignment using Expresso showed EcoNei has a 7 amino acid insertion in the linker region compared to human NEIL1 and mimivirus Nei1 (71). There were no notable differences in linker size between the FPG enzymes, although this conformational change is much more modest compared to EcoNei and hNTHL1. With the accumulation of structural evidence, interdomain movements of the DNA glycosylases may in fact be a more common mechanism than previously thought, albeit with differing degrees of movement.

Investigations of protein movement in solution with tryptophan fluorescence have shown that EcoNei, EcoFpg, human NEIL1, and human OGG1 undergo protein isomerization events prior to catalysis, but the degree of movement is unknown because tryptophan fluorescence simply indicates that the environment of the tryptophan has changed (68,72–74). Changes in fluorescence were not detected for EcoNth, suggesting that either EcoNth does not undergo a conformational change during catalysis, or that it was simply not observed under the studied conditions because of the non-optimal location of tryptophan residues within the protein (75). Our tryptophan fluorescence data yielded a large difference in signal between hNTHL1Δ63 and hNTHL1Δ63 chimera in the presence of Tg:A - containing DNA, but not with THF:A or undamaged DNA. This finding suggests that there is a large conformational change at some point after lesion recognition, as the undamaged and THF curves look similar for NTHL1Δ63.

In addition to its interdomain flexibility, hNTHL1 also differs from its prokaryotic homologs because it possesses a flexible N-terminal extension (residues 1 to 86) (Figure 1C). This N-terminal extension was posited to inhibit product release when hNTHL1 is at low concentration (40). The emergence of a sigmoidal curve when DNA glycosylase activity is plotted *vs*. enzyme concentration hinted at the possibility of positive cooperativity, leading to the hypothesis that hNTHL1 forms dimers (39,41). A crosslinking experiment with BS3, an amine-to-amine crosslinker, showed that serial truncations of the N-terminal extension reduced the amount of crosslinked hNTHL1 and putative dimers (39). However, we propose that the decreased propensity to form dimers as the N-terminus was progressively trimmed is likely to be due to the serial elimination of crosslinkable lysines. We sought to identify the predicted hNTHL1 dimer, but the protein consistently eluted off the sizing column (Superdex 75; Cytiva) as a monomer (Figure S7). Additionally, analysis of the crystal lattice of hNTHL1Δ63, which still maintains 27 residues of the N-terminal extension, did not reveal any putative dimerization interface.

How, then, could NTHL1 display positive cooperativity in the absence of dimer formation? Another means to achieve apparent enzymatic cooperativity independent of multimerization is kinetic cooperativity (76). A well-documented instance of kinetic cooperativity (or monomeric enzyme cooperativity) is glucokinase, which exemplifies this particular behaviour in both kinetic and structural data (76). Glucokinase performs the first step in glycolysis by phosphorylation of glucose to glucose-6-phosphate. Glucokinase comprises two domains, and the active site is formed in the cleft between those domains. Much like hNTHL1, the glucokinase structures revealed a hinge region, with a 99° rotation of the two domains between the open and closed conformations (77). When quantifying the biochemical reaction trajectories, glucokinase exhibits positive cooperativity, evident as a sigmoidal curve relating substrate concentration to rate, in the presence of increasing glucose concentration (78). Kamata *et al*. interpreted the crystal structures, postulating the existence of a “super-open” ground state conformation that is driven to an open glucose-bound state. This equilibrium depends on the concentration of glucose, and is rate-limiting in the greater reaction scheme. Sigmoidal reaction curves therefore emerge due to bypass of this rate-limiting step at high concentrations of glucose, rather than by the conventional cooperative model in which the affinity of the binding sites for a ligand is increased.

In the context of our current understanding of hNTHL1, we speculate that the DNA glycosylase forms an open scanning configuration that can slide along the DNA searching for a lesion. Upon encountering a lesion, hNTHL1 would close on the DNA to process the lesion. After processing, the enzyme would release back into the scanning complex to continue its search for lesions (Figure 5). This model is supported by the two conformations of hNTHL1, in open and closed states. Furthermore, the tryptophan fluorescence stopped-flow data are consistent with this hypothesis. hNTHL1 exhibits a low affinity for undamaged DNA (30 μM (79) *vs*. nM range for lesion-containing DNA (19)), therefore we can assume that there is little appreciable DNA binding signal in the undamaged DNA curve. hNTHL1 binds to DNA containing THF but, as this lesion is non-cleavable, the enzyme cannot cycle through catalysis, thus keeping the enzyme in a lesion-bound state. In contrast, hNTHL1 can bind, cleave, and release Tg, and this is the only DNA substrate for which a change in signal is observed. This finding suggests that a conformational change occurs either during or after the cleavage event, perhaps to protect undamaged bases from being erroneously cleaved. It is important to note that tryptophan fluorescence relies on a change in environment around a particular tryptophan (Figure S5), and therefore we cannot rule out the possibility of additional conformational changes during the catalytic cycle.

**Figure 5:**
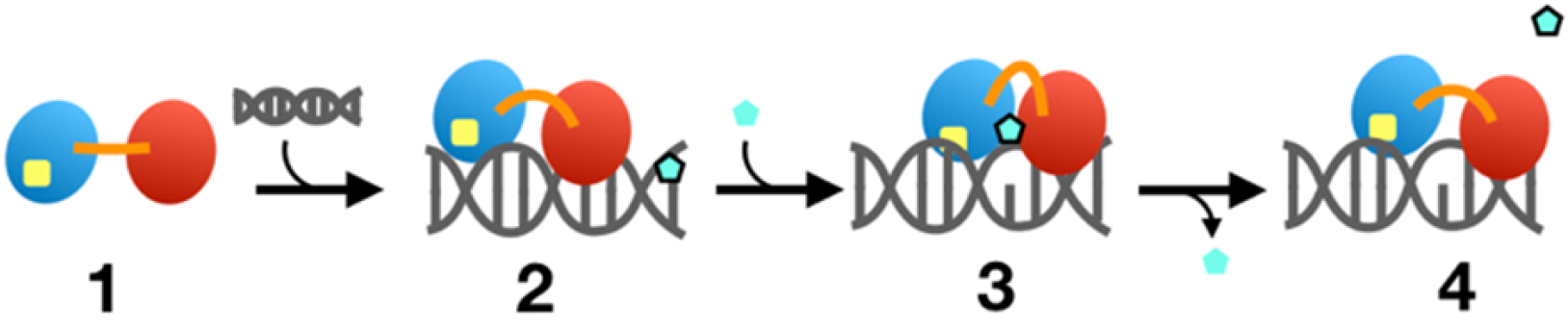
Model for NTHL1 conformational change. Proposed model of hNTHL1 lesion searching, recognition and removal, which is consistent with kinetic cooperativity. The [4Fe4S] domain is shown in blue, with the [4Fe4S] cluster represented in yellow; the six-helical bundle domain is red and the interdomain flexible linker region is orange. In Step 1, the apo enzyme exists in an open conformation; when it associates with undamaged DNA, the DNA glycosylase adopts a semi-closed state, or “scanning conformation”, depicted in Step 2. As hNTHL1 scans the DNA, if it encounters a lesion (cyan pentagon) the closed or “active” complex is achieved, Step 3. After lesion removal the enzyme may relax into the scanning conformation to continue a search for DNA damage (Step 4).

### Conclusion

The open conformation observed in hNTHL1 presents a novel molecular mechanism for this DNA glycosylase. The bacterial homologs that laid the foundation for the current kinetics models do not appear to undergo a conformational change, based on the available X-ray crystal structures and solution fluorescence studies (20,21,56,75). We have established that a truncation of linker 1 in hNTHL1 shifts the equilibrium towards the closed conformation. The reduced interdomain flexibility of hNTHL1Δ63 chimera decreased the binding affinity to DNA. The hNTHL1Δ63 chimera, however, still retains glycosylase and β-lyase activity, indicating that the active site is properly formed. Interdomain movements have been observed in the H2TH and UDG families of glycosylases, and now the HhH family, indicating that this mechanism may be more common than previously thought for DNA glycosylases. Analysis of previously reported variant data and germline SNVs suggests that the conformational change of hNTHL1 is crucial for proper function.

## ACCESSION NUMBERS

Atomic coordinates and structure factors for the reported crystal structures have been deposited with the Protein Data Bank (www.rcsb.org) under accession numbers 7RDS (hNTHL1Δ63) and 7RDT (hNTHL1Δ63 chimera).

## SUPPLEMENTARY DATA

Supplementary Data are available at NAR online.

## ACKNOWLEDGEMENTS

We thank Drs. Joann Sweasy and Khadijeh Alnajjar for help with the stopped-flow SX-20 instrument, Dr. Brian Eckenroth for crystallographic data collection of the hNTHL1Δ63 chimera, and Dr. Rick Wood for providing comments on the manuscript. Identification and selection of human variants along with DNA sequencing services were performed using the Bioinformatics Shared Resource and Vermont Integrative Genomics Resource at the University of Vermont. The germline SNVs shown are based upon data generated by the TCGA Research Network https://www.cancer.gov/tcga. Support of the X-ray facility by the Trunk Foundation is gratefully acknowledged.

---

**Table S1:** Germline hNTHL1 variants from the TCGA dataset. The germline SNVs curated from the TCGA dataset are listed in ascending order. The RSID number is listed, if available, as well as the SNV frequency and REVEL (55) score.

| RSIDnumber | Missense | SNV Frequency | REVEL |
| --- | --- | --- | --- |
| rs148104494 | R100C | 16 | 0.744 |
| rs148104494 | R100S | 4 | 0.735 |
|  | G114V | 5 | 0.788 |
|  | G114W | 16 | 0.849 |
|  | P125L | 1 | 0.579 |
|  | P125Q | 8 | 0.708 |
| rs149277519 | P125S | 1 | 0.627 |
| rs149277519 | P125T | 5 | 0.653 |
| rs1805378 | I176T | 36 | 0.876 |
| rs759754271 | I204T | 30 | 0.635 |
|  | V228G | 41 | 0.686 |
|  | W230R | 15 | 0.895 |
|  | W251G | 9 | 0.762 |
|  | W251R | 19 | 0.741 |
| rs1437503034 | P259Q | 9 | 0.541 |
| rs367765503 | P259T | 12 | 0.512 |
| rs779992803 | R263C | 3 | 0.517 |
| rs779992803 | R263S | 29 | 0.558 |
| rs148474733 | A265P | 1 | 0.606 |
|  | P271A | 349 | 0.759 |
|  | T289P | 6 | 0.617 |

**Figure S1:**
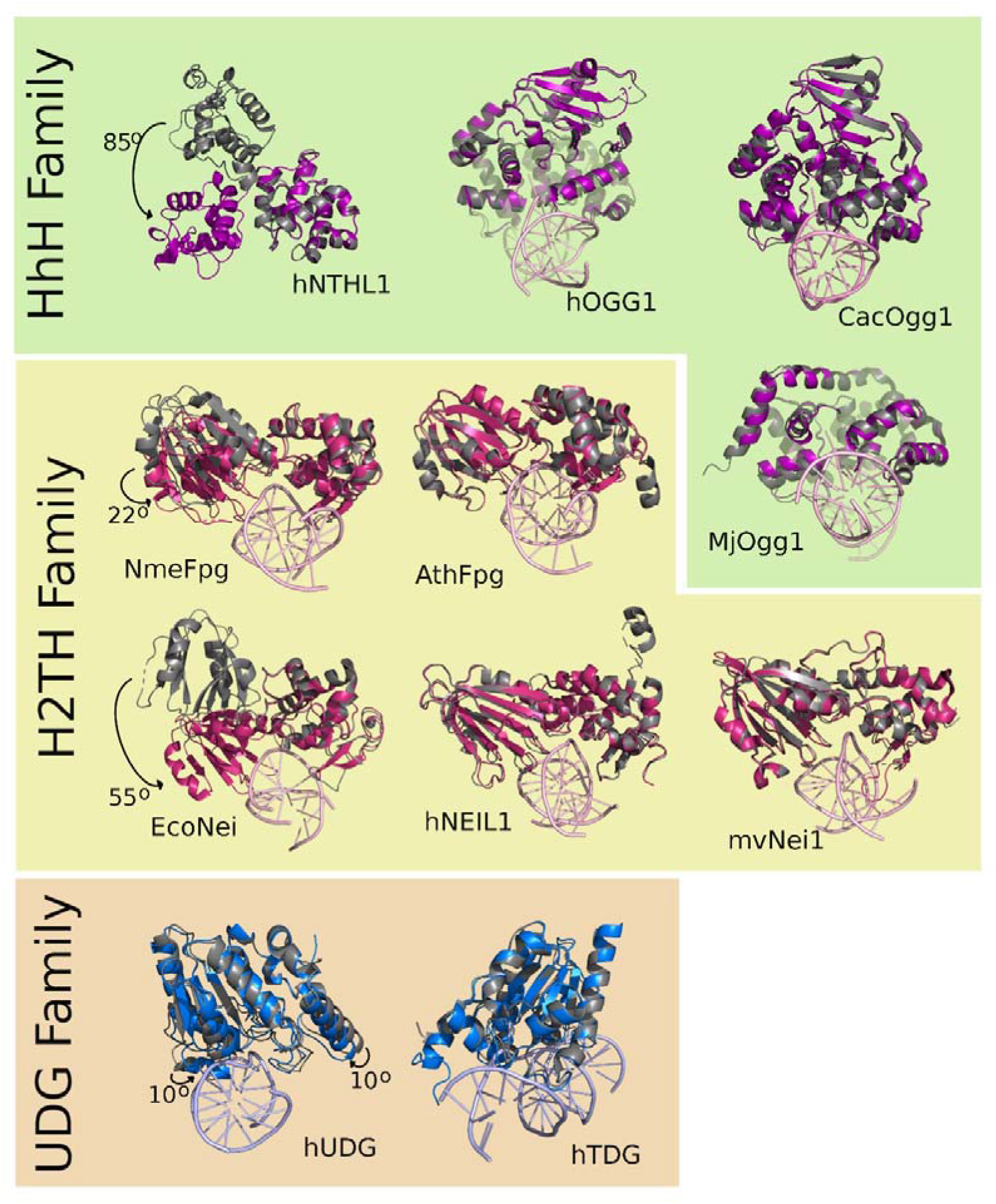
Domain orientation of DNA glycosylases crystallized with and without DNA. A compilation of liganded and unliganded DNA glycosylase crystal structures reveals a few DNA glycosylases with gross interdomain movements. In the UDG family, human UDG has a 10° “pinching” in of the open, unbound conformation (PDB ID code 1AKZ, grey) to meet the closed, bound conformation (2SSP, blue) (70). In the H2TH family, EcoNei exhibits an interdomain rotation of 50° between the apo (1Q39, grey) and bound forms (1K3W, red) (63,68), and NmeFPG has a 22° rotation between unbound (6TC6, grey) and bound (6TC9, red) (81). In the HhH family, hNTHL1 undergoes an 85° rotation between open (7RDS, grey) and closed conformations (7RDT, purple). In contrast, other DNA glycosylases exhibit near identical conformations between unliganded and DNA-bound states: human TDG unbound (1WYW, grey) and DNA-bound (5T2W, blue) (82,83); human NEIL1 apo (1TDH, grey) and DNA-bound (5ITR, red) (84,85), *Mimivirus* Nei1 apo (3A42, grey) and DNA-bound (3A46, red) (86), *Arabidopsis thaliana* Fpg apo (3TWL, grey) and DNA-bound (3TWM, red) (87); human OGG1 unbound (1KO9, grey) and bound (1EBM, purple) (88,89), *Methanocaldococcus jannaschii* Ogg unbound (3FHF, grey) and bound (3KNT, purple) (90,91), *Clostridium acetobutylicum* Ogg1 unbound (3F0Z, grey) and bound (3I0X, purple) (92,93).

**Figure S2:**
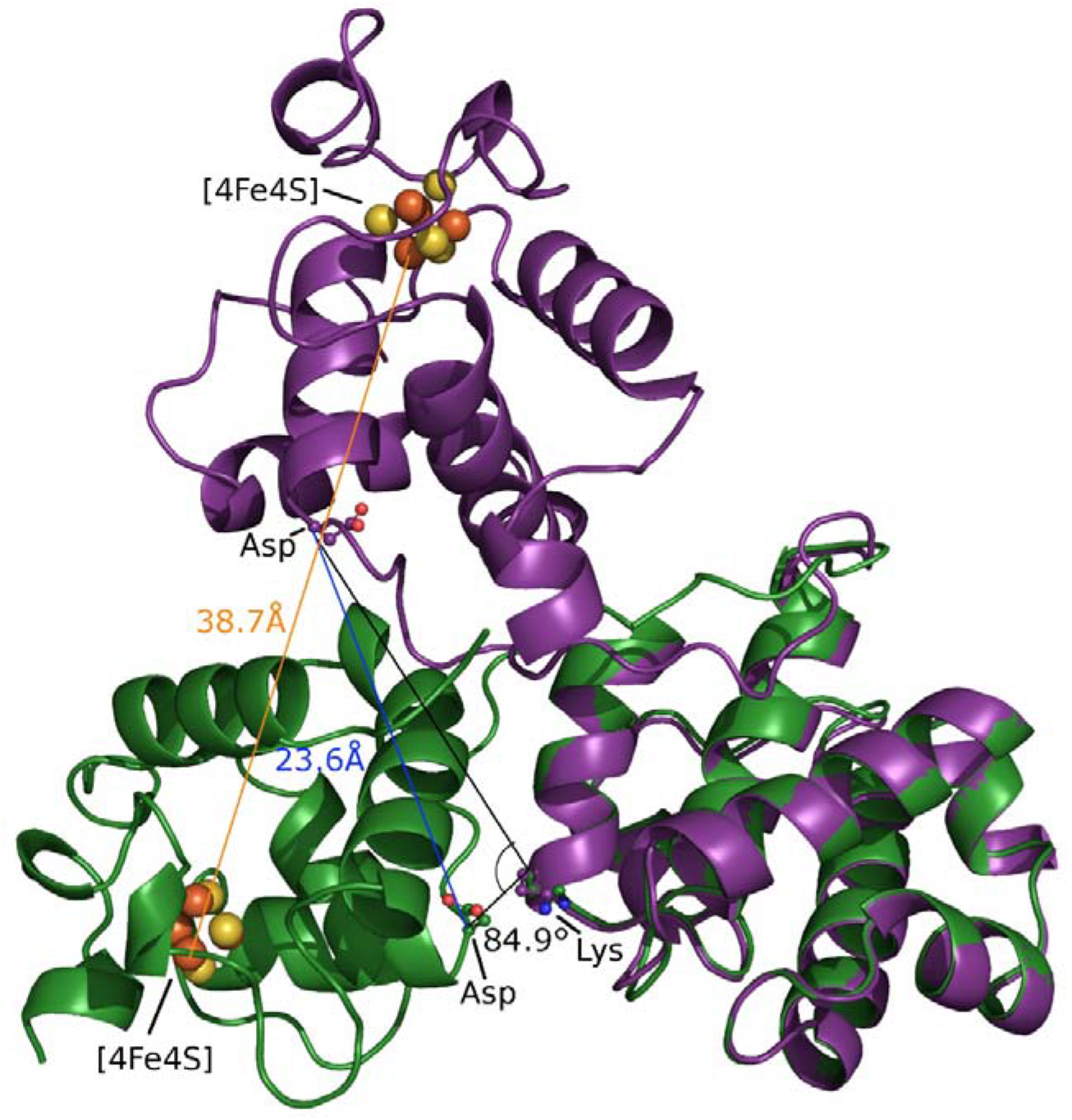
hNTHL1Δ63 and hNTHL1Δ63 chimera overlay. This figure illustrates the measurements made between hNTHL1Δ63 (purple) and hNTHL1Δ63 chimera (green). The superposition between the two models was performed with PyMOL using the 6-helical bundle domain as reference (59).

**Figure S3:**
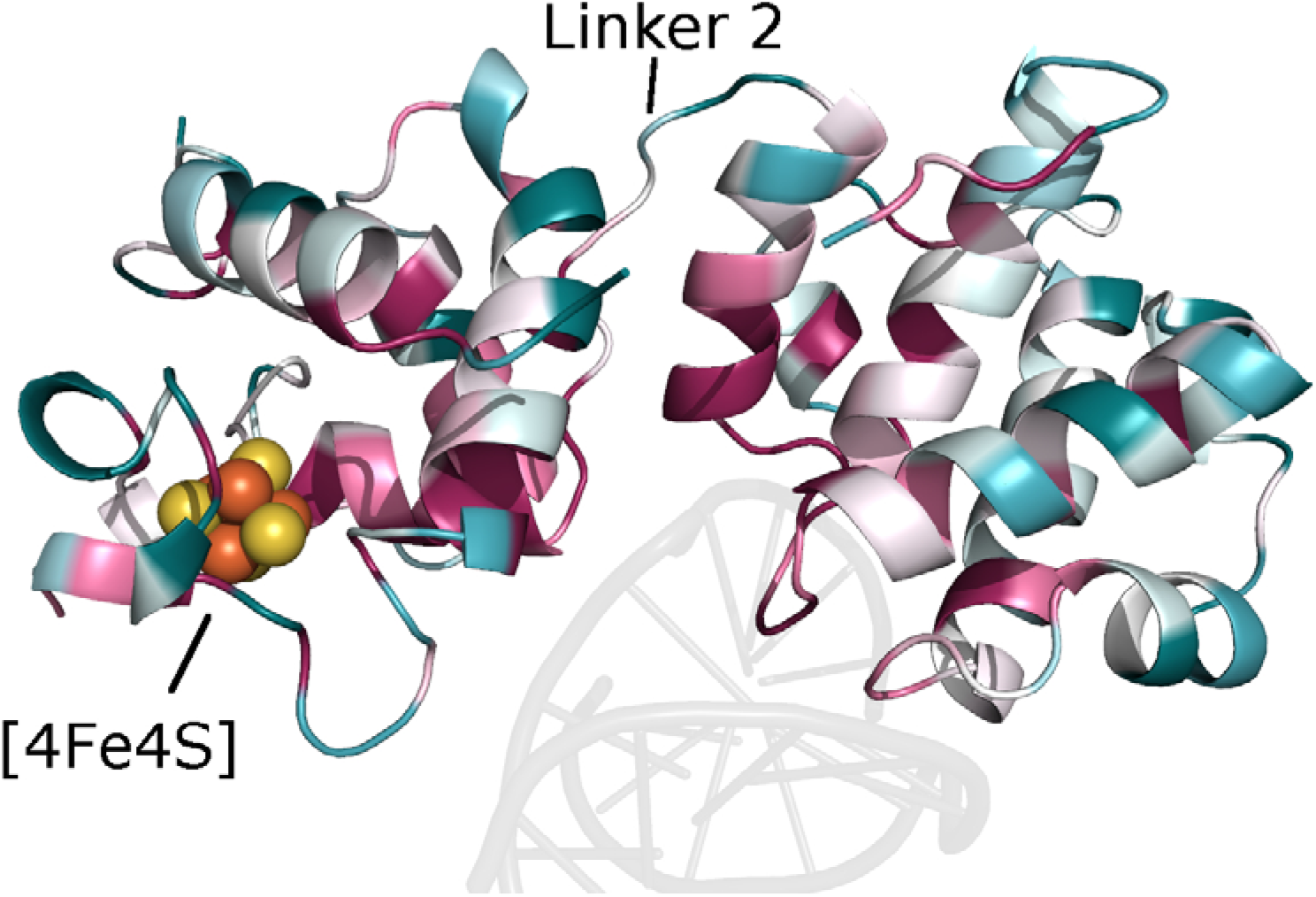
The DNA binding cleft is conserved. The NTHL1Δ63 chimera model coloured according to amino acid conservation (red conserved, blue variable) using Consurf (65) shows that the DNA binding regions are conserved.

**Figure S4:**
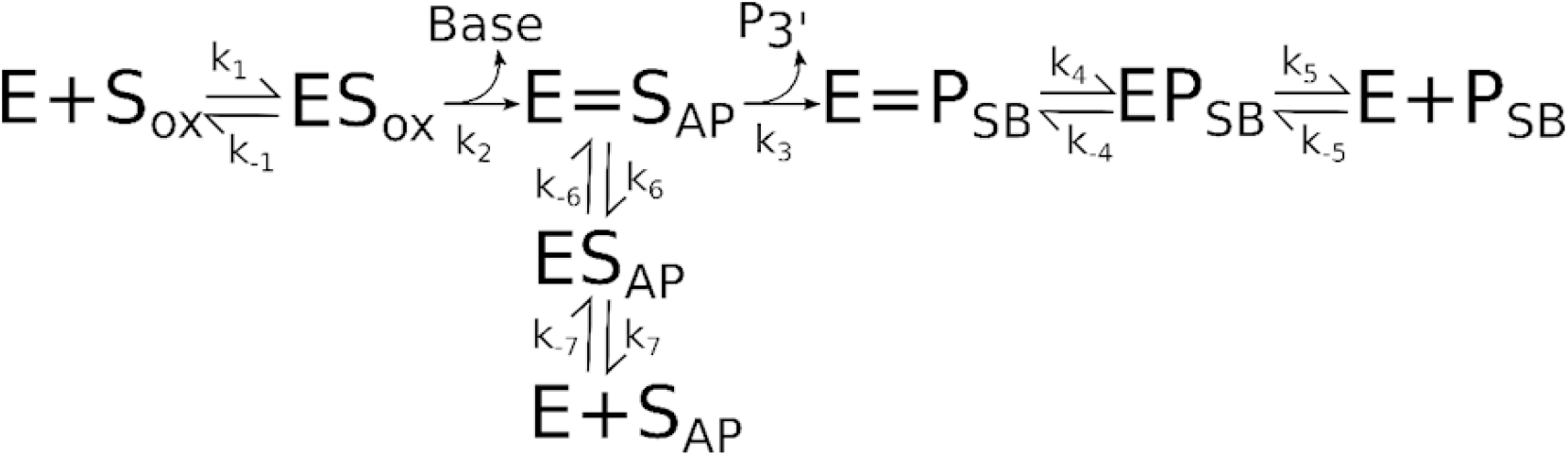
hNTHL1 schematic of enzymatic steps based upon (60). hNTHL1 (E) binds the oxidized DNA lesion (S_OX_), upon N-glycosidic cleavage the base is released, and the substrate is now an abasic site (S_AP_). During the breakage of the N-glycosidic bond a Schiff base is created between the enzyme and substrate. hNTHL1 is bifunctional, therefore the β-lytic activity releases the 3’-end of the β-eliminated deoxynucleotide (P_3′_). After β-elimination, resolution of the Schiff base precedes the single strand break product release (P_SB_). If-elimination does not occur, then the Schiff base is resolved from the abasic site, and the final product is S_AP_. The rate constants k_1_ and k_-1_ are the forward and reverse rates of substrate association and dissociation, k_2_ is the rate of base excision or glycosylase activity, k_3_ is the rate of β-lyase activity, k_4_ and k_-4_ are the forward and reverse rates of Schiff base resolution, k_5_ and k_-5_ are the forward and reverse rates of product (single strand break) release, k_6_ and k_-6_ are the forward and reverse rates of Schiff base resolution without β-lyase occurring prior, and k_7_ and k_-7_ are the forward and reverse rates of AP site release or re-association.

**Figure S5:**
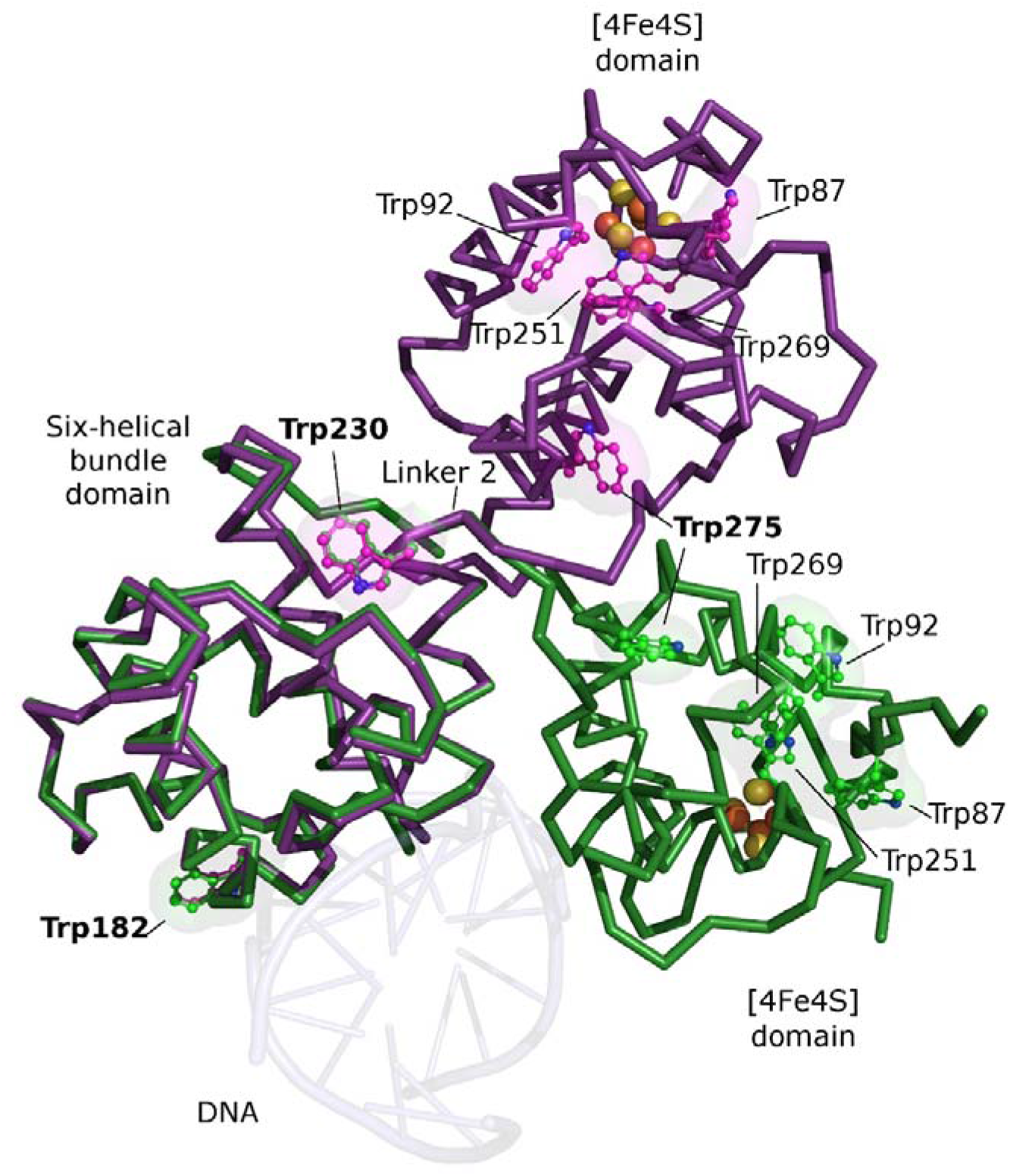
Mapping of tryptophan residues onto the hNTHL1Δ63 and hNTHL1Δ63 chimera models. This figure shows the location of the seven tryptophan residues in hNTHL1. Four tryptophans are clustered together near the [4Fe4S] cluster, two are located near linker 2, and one is in the six-helical bundle domain. In this figure the six-helical bundle domain is shown on the left to better highlight the position of all tryptophan residues in hNTHL1.

**Figure S6:**
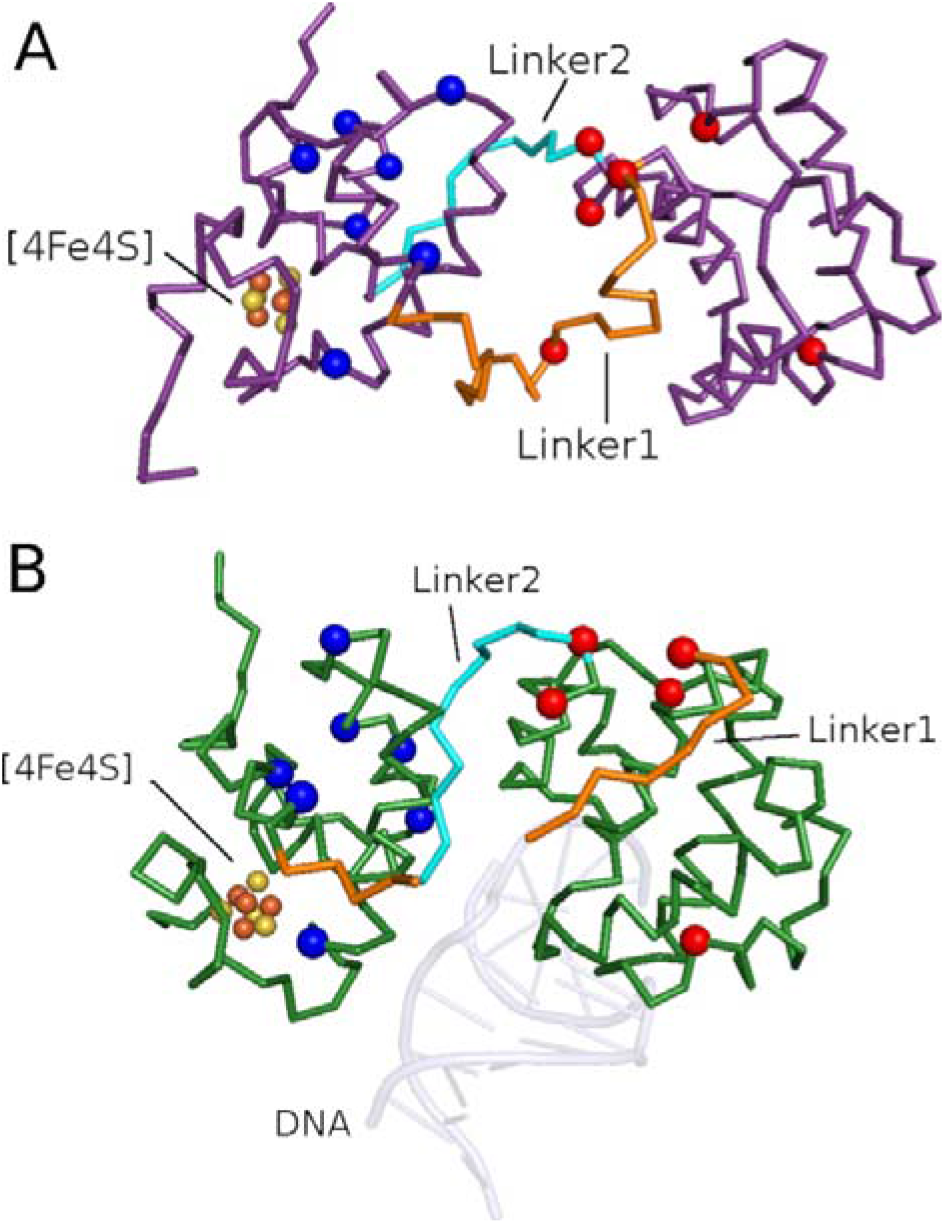
Mapping of cancer variants onto the hNTHL1Δ63 closed structure. Germline variants are grouped in clusters: SNVs are found within the six-helical bundle: Gly114Val/T rp, Pro125Leu/Gln/Ser/Thr, Ile176Thr, Ile204Thr, Val228Gly, and Trp230Arg (shown as red spheres), and the [4Fe4S] domain: Arg100Cys/Ser, Trp251Gly/Arg, Pro259Gln/Thr, Arg263Cys/Ser, Ala265Pro, Pro271Ala, and Thr289Pro (blue spheres) in A) the open hNTHL1 model and B) the closed hNTHL1 chimera model with DNA from GstNth (PDB 1ORN, (21)). The variants cluster away from the DNA binding interface, and in the vicinity of the two interdomain linker regions.

**Figure S7:**
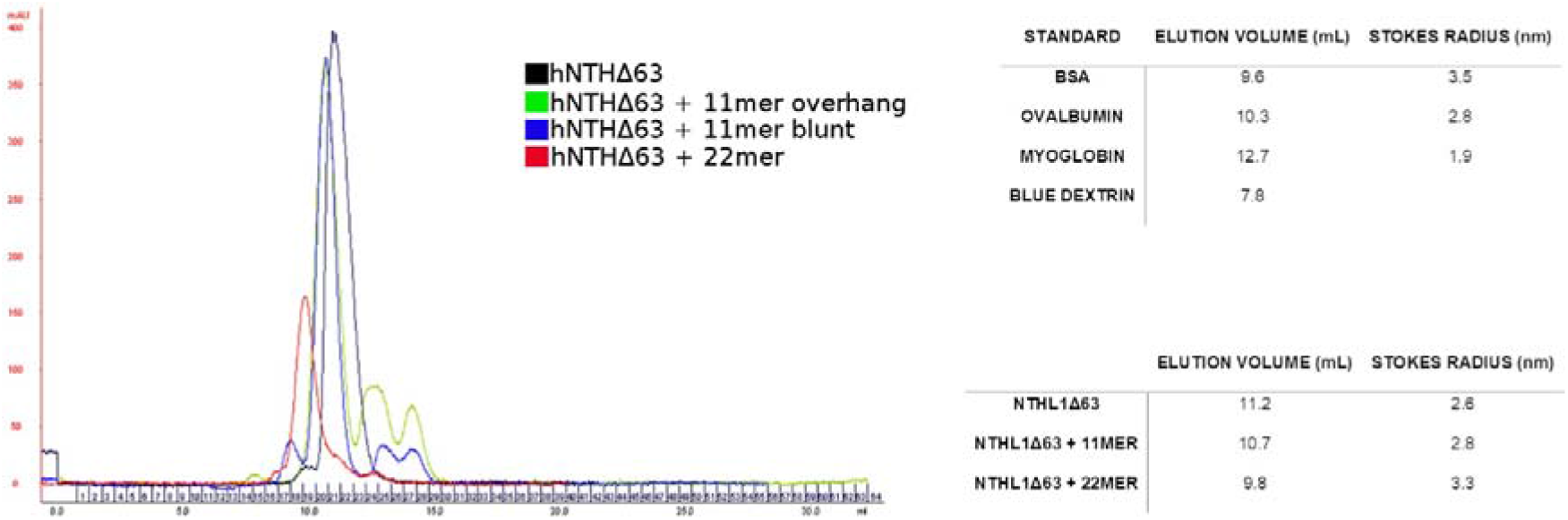
Elution profile of hNTHL1Δ63 from a gel filtration column. The elution profiles of hNTHL1Δ63 with and without DNA substrates on a Superdex 75 column (Cytiva) are shown. The Stokes radius of 2.6 nm fits that of a monomer of hNTHL1Δ63. The radius of gyration calculated with PyMOL for hNTHL1Δ63 is 2.1 nm (59). We used two different lengths of DNA: the 11mer should bind a single monomer, and the 22mer should be able to accommodate two molecules. In all cases the predominant species appears to be a monomer. Equation for line of best fit: −1.9277x+16.132 R^2^=0.9437.

## REFERENCES

1. Wallace, S.S. (2014) Base excision repair: a critical player in many games. DNA Repair (Amst), 19, 14–26.

2. Brooks, S.C., Adhikary, S., Rubinson, E.H. and Eichman, B.F. (2013) Recent advances in the structural mechanisms of DNA glycosylases. Biochim Biophys Acta, 1834, 247–271.

3. Bauer, N.C., Corbett, A.H. and Doetsch, P.W. (2015) The current state of eukaryotic DNA base damage and repair. Nucleic Acids Res, 43, 10083–10101.

4. Prasad, R., Shock, D.D., Beard, W.A. and Wilson, S.H. (2010) Substrate channeling in mammalian base excision repair pathways: passing the baton. J Biol Chem, 285, 40479–40488.

5. Wilson, S.H. and Kunkel, T.A. (2000) Passing the baton in base excision repair. Nat Struct Biol, 7, 176–178.

6. Dianov, G., Bischoff, C., Piotrowski, J. and Bohr, V.A. (1998) Repair pathways for processing of 8-oxoguanine in DNA by mammalian cell extracts. J Biol Chem, 273, 33811–33816.

7. Fortini, P., Parlanti, E., Sidorkina, O.M., Laval, J. and Dogliotti, E. (1999) The type of DNA glycosylase determines the base excision repair pathway in mammalian cells. J Biol Chem, 274, 15230–15236.

8. Cappelli, E., Taylor, R., Cevasco, M., Abbondandolo, A., Caldecott, K. and Frosina, G. (1997) Involvement of XRCC1 and DNA ligase III gene products in DNA base excision repair. J Biol Chem, 272, 23970–23975.

9. Mol, C.D., Izumi, T., Mitra, S. and Tainer, J.A. (2000) DNA-bound structures and mutants reveal abasic DNA binding by APE1 and DNA repair coordination. Nature, 403, 451–456.

10. Waters, T.R., Gallinari, P., Jiricny, J. and Swann, P.F. (1999) Human thymine DNA glycosylase binds to apurinic sites in DNA but is displaced by human apurinic endonuclease 1. J Biol Chem, 274, 67–74.

11. Vidal, A.E., Hickson, I.D., Boiteux, S. and Radicella, J.P. (2001) Mechanism of stimulation of the DNA glycosylase activity of hOGG1 by the major human AP endonuclease: bypass of the AP lyase activity step. Nucleic Acids Res, 29, 1285–1292.

12. Hill, J.W., Hazra, T.K., Izumi, T. and Mitra, S. (2001) Stimulation of human 8-oxoguanine-DNA glycosylase by AP-endonuclease: potential coordination of the initial steps in base excision repair. Nucleic Acids Res, 29, 430–438.

13. Maher, R.L., Wallace, S.S. and Pederson, D.S. (2019) The lyase activity of bifunctional DNA glycosylases and the 3’-diesterase activity of APE1 contribute to the repair of oxidized bases in nucleosomes. Nucleic Acids Res, 47, 2922–2931.

14. Boiteux, S. and Guillet, M. (2004) Abasic sites in DNA: repair and biological consequences in Saccharomyces cerevisiae. DNA Repair (Amst), 3, 1–12.

15. Simonelli, V., Narciso, L., Dogliotti, E. and Fortini, P. (2005) Base excision repair intermediates are mutagenic in mammalian cells. Nucleic Acids Res, 33, 4404–4411.

16. Katcher, H.L. and Wallace, S.S. (1983) Characterization of the Escherichia coli X-ray endonuclease, endonuclease III. Biochemistry, 22, 4071–4081.

17. Aspinwall, R., Rothwell, D.G., Roldan-Arjona, T., Anselmino, C., Ward, C.J., Cheadle, J.P., Sampson, J.R., Lindahl, T., Harris, P.C. and Hickson, I.D. (1997) Cloning and characterization of a functional human homolog of Escherichia coli endonuclease III. Proc Natl Acad Sci U S A, 94, 109–114.

18. Ikeda, S., Biswas, T., Roy, R., Izumi, T., Boldogh, I., Kurosky, A., Sarker, A.H., Seki, S. and Mitra, S. (1998) Purification and characterization of human NtH1, a homolog of Escherichia coli endonuclease III. Direct identification of Lys-212 as the active nucleophilic residue. J Biol Chem, 273, 21585–21593.

19. Robey-Bond, S.M., Benson, M.A., Barrantes-Reynolds, R., Bond, J.P. and Wallace, S.S. (2017) Probing the activity of NTHL1 orthologs by targeting conserved amino acid residues. DNA Repair (Amst), 53, 43–51.

20. Kuo, C.F., McRee, D.E., Fisher, C.L., O’Handley, S.F., Cunningham, R.P. and Tainer, J.A. (1992) Atomic structure of the DNA repair [4Fe-4S] enzyme endonuclease III. Science, 258, 434–440.

21. Fromme, J.C. and Verdine, G.L. (2003) Structure of a trapped endonuclease III-DNA covalent intermediate. EMBO J, 22, 3461–3471.

22. Wallace, S.S., Murphy, D.L. and Sweasy, J.B. (2012) Base excision repair and cancer. Cancer Lett, 327, 73–89.

23. Das, L., Quintana, V.G. and Sweasy, J.B. (2020) NTHL1 in genomic integrity, aging and cancer. DNA Repair (Amst), 93, 102920.

24. Weren, R.D., Ligtenberg, M.J., Kets, C.M., de Voer, R.M., Verwiel, E.T., Spruijt, L., van Zelst-Stams, W.A., Jongmans, M.C., Gilissen, C., Hehir-Kwa, J.Y. et al. (2015) A germline homozygous mutation in the base-excision repair gene NTHL1 causes adenomatous polyposis and colorectal cancer. Nat Genet, 47, 668–671.

25. Valle, L., de Voer, R.M., Goldberg, Y., Sjursen, W., Forsti, A., Ruiz-Ponte, C., Caldes, T., Garre, P., Olsen, M.F., Nordling, M. et al. (2019) Update on genetic predisposition to colorectal cancer and polyposis. Mol Aspects Med, 69, 10–26.

26. Broderick, P., Dobbins, S.E., Chubb, D., Kinnersley, B., Dunlop, M.G., Tomlinson, I. and Houlston, R.S. (2017) Validation of Recently Proposed Colorectal Cancer Susceptibility Gene Variants in an Analysis of Families and Patients-a Systematic Review. Gastroenterology, 152, 75–77 e74.

27. Fostira, F., Kontopodis, E., Apostolou, P., Fragkaki, M., Androulakis, N., Yannoukakos, D., Konstantopoulou, I. and Saloustros, E. (2018) Extending the clinical phenotype associated with biallelic NTHL1 germline mutations. Clin Genet, 94, 588–589.

28. Grolleman, J.E., de Voer, R.M., Elsayed, F.A., Nielsen, M., Weren, R.D.A., Palles, C., Ligtenberg, M.J.L., Vos, J.R., Ten Broeke, S.W., de Miranda, N. et al. (2019) Mutational Signature Analysis Reveals NTHL1 Deficiency to Cause a Multi-tumor Phenotype. Cancer Cell, 35, 256–266 e255.

29. Groves, A., Gleeson, M. and Spigelman, A.D. (2019) NTHL1-associate polyposis: first Australian case report. Fam Cancer, 18, 179–182.

30. Rivera, B., Castellsague, E., Bah, I., van Kempen, L.C. and Foulkes, W.D. (2015) Biallelic NTHL1 Mutations in a Woman with Multiple Primary Tumors. N Engl J Med, 373, 1985–1986.

31. Belhadj, S., Mur, P., Navarro, M., Gonzalez, S., Moreno, V., Capella, G. and Valle, L. (2017) Delineating the Phenotypic Spectrum of the NTHL1-Associated Polyposis. Clin Gastroenterol Hepatol, 15, 461–462.

32. Weren, R.D., Ligtenberg, M.J., Geurts van Kessel, A., De Voer, R.M., Hoogerbrugge, N. and Kuiper, R.P. (2018) NTHL1 and MUTYH polyposis syndromes: two sides of the same coin? J Pathol, 244, 135–142.

33. Belhadj, S., Quintana, I., Mur, P., Munoz-Torres, P.M., Alonso, M.H., Navarro, M., Terradas, M., Pinol, V., Brunet, J., Moreno, V. et al. (2019) NTHL1 biallelic mutations seldom cause colorectal cancer, serrated polyposis or a multi-tumor phenotype, in absence of colorectal adenomas. Sci Rep, 9, 9020.

34. Terradas, M., Munoz-Torres, P.M., Belhadj, S., Aiza, G., Navarro, M., Brunet, J., Capella, G. and Valle, L. (2019) Contribution to colonic polyposis of recently proposed predisposing genes and assessment of the prevalence of NTHL1- and MSH3-associated polyposes. Hum Mutat, 40, 1910–1923.

35. Drost, J., van Boxtel, R., Blokzijl, F., Mizutani, T., Sasaki, N., Sasselli, V., de Ligt, J., Behjati, S., Grolleman, J.E., van Wezel, T. et al. (2017) Use of CRISPR-modified human stem cell organoids to study the origin of mutational signatures in cancer. Science, 358, 234–238.

36. Nik-Zainal, S., Davies, H., Staaf, J., Ramakrishna, M., Glodzik, D., Zou, X., Martincorena, I., Alexandrov, L.B., Martin, S., Wedge, D.C. et al. (2016) Landscape of somatic mutations in 560 breast cancer whole-genome sequences. Nature, 534, 47–54.

37. Wong, H.L., Yang, K.C., Shen, Y., Zhao, E.Y., Loree, J.M., Kennecke, H.F., Kalloger, S.E., Karasinska, J.M., Lim, H.J., Mungall, A.J. et al. (2018) Molecular characterization of metastatic pancreatic neuroendocrine tumors (PNETs) using whole-genome and transcriptome sequencing. Cold Spring Harb Mol Case Stud, 4.

38. Limpose, K.L., Trego, K.S., Li, Z., Leung, S.W., Sarker, A.H., Shah, J.A., Ramalingam, S.S., Werner, E.M., Dynan, W.S., Cooper, P.K. et al. (2018) Overexpression of the base excision repair NTHL1 glycosylase causes genomic instability and early cellular hallmarks of cancer. Nucleic Acids Res, 46, 4515–4532.

39. Liu, X., Choudhury, S. and Roy, R. (2003) In vitro and in vivo dimerization of human endonuclease III stimulates its activity. J Biol Chem, 278, 50061–50069.

40. Liu, X. and Roy, R. (2002) Truncation of amino-terminal tail stimulates activity of human endonuclease III (hNTH1). J Mol Biol, 321, 265–276.

41. Marenstein, D.R., Ocampo, M.T., Chan, M.K., Altamirano, A., Basu, A.K., Boorstein, R.J., Cunningham, R.P. and Teebor, G.W. (2001) Stimulation of human endonuclease III by Y box-binding protein 1 (DNA-binding protein B). Interaction between a base excision repair enzyme and a transcription factor. J Biol Chem, 276, 21242–21249.

42. Studier, F.W. (2005) Protein production by auto-induction in high density shaking cultures. Protein Expr Purif, 41, 207–234.

43. Doublie, S. (2007) Production of selenomethionyl proteins in prokaryotic and eukaryotic expression systems. Methods Mol Biol, 363, 91–108.

44. Evans, P.R. (2011) An introduction to data reduction: space-group determination, scaling and intensity statistics. Acta Crystallogr D Biol Crystallogr, 67, 282–292.

45. Evans, P.R. and Murshudov, G.N. (2013) How good are my data and what is the resolution? Acta Crystallogr D Biol Crystallogr, 69, 1204–1214.

46. Evans, P. (2006) Scaling and assessment of data quality. Acta Crystallogr D Biol Crystallogr, 62, 72–82.

47. Winn, M.D., Ballard, C.C., Cowtan, K.D., Dodson, E.J., Emsley, P., Evans, P.R., Keegan, R.M., Krissinel, E.B., Leslie, A.G., McCoy, A. et al. (2011) Overview of the CCP4 suite and current developments. Acta Crystallogr D Biol Crystallogr, 67, 235–242.

48. Battye, T.G., Kontogiannis, L., Johnson, O., Powell, H.R. and Leslie, A.G. (2011) iMOSFLM: a new graphical interface for diffraction-image processing with MOSFLM. Acta Crystallogr D Biol Crystallogr, 67, 271–281.

49. Adams, P.D., Afonine, P.V., Bunkoczi, G., Chen, V.B., Davis, I.W., Echols, N., Headd, J.J., Hung, L.W., Kapral, G.J., Grosse-Kunstleve, R.W. et al. (2010) PHENIX: a comprehensive Python-based system for macromolecular structure solution. Acta Crystallogr D Biol Crystallogr, 66, 213–221.

50. Terwilliger, T.C., Read, R.J., Adams, P.D., Brunger, A.T., Afonine, P.V. and Hung, L.W. (2013) Model morphing and sequence assignment after molecular replacement. Acta Crystallogr D Biol Crystallogr, 69, 2244–2250.

51. Afonine, P.V., Grosse-Kunstleve, R.W., Urzhumtsev, A. and Adams, P.D. (2009) Automatic multiple-zone rigid-body refinement with a large convergence radius. J Appl Crystallogr, 42, 607–615.

52. Afonine, P.V., Grosse-Kunstleve, R.W., Adams, P.D. and Urzhumtsev, A. (2013) Bulk-solvent and overall scaling revisited: faster calculations, improved results. Acta Crystallogr D Biol Crystallogr, 69, 625–634.

53. Headd, J.J., Echols, N., Afonine, P.V., Grosse-Kunstleve, R.W., Chen, V.B., Moriarty, N.W., Richardson, D.C., Richardson, J.S. and Adams, P.D. (2012) Use of knowledge-based restraints in phenix.refine to improve macromolecular refinement at low resolution. Acta Crystallogr D Biol Crystallogr, 68, 381–390.

54. Afonine, P.V., Grosse-Kunstleve, R.W., Echols, N., Headd, J.J., Moriarty, N.W., Mustyakimov, M., Terwilliger, T.C., Urzhumtsev, A., Zwart, P.H. and Adams, P.D. (2012) Towards automated crystallographic structure refinement with phenix.refine. Acta Crystallogr D Biol Crystallogr, 68, 352–367.

55. Ioannidis, N.M., Rothstein, J.H., Pejaver, V., Middha, S., McDonnell, S.K., Baheti, S., Musolf, A., Li, Q., Holzinger, E., Karyadi, D. et al. (2016) REVEL: An Ensemble Method for Predicting the Pathogenicity of Rare Missense Variants. Am J Hum Genet, 99, 877–885.

56. Cunningham, R.P., Ahern, H., Xing, D., Thayer, M.M. and Tainer, J.A. (1994) Structure and function of Escherichia coli endonuclease III. Ann N Y Acad Sci, 726, 215–222.

57. Sarre, A., Stelter, M., Rollo, F., De Bonis, S., Seck, A., Hognon, C., Ravanat, J.L., Monari, A., Dehez, F., Moe, E. et al. (2019) The three Endonuclease III variants of Deinococcus radiodurans possess distinct and complementary DNA repair activities. DNA Repair (Amst), 78, 45–59.

58. Thayer, M.M., Ahern, H., Xing, D., Cunningham, R.P. and Tainer, J.A. (1995) Novel DNA binding motifs in the DNA repair enzyme endonuclease III crystal structure. EMBO J, 14, 4108–4120.

59. The Pymol Molecular Graphics System, Version 1.2r3pre, Schrodinger, LLC.

60. Galick, H.A., Kathe, S., Liu, M., Robey-Bond, S., Kidane, D., Wallace, S.S. and Sweasy, J.B. (2013) Germ-line variant of human NTH1 DNA glycosylase induces genomic instability and cellular transformation. Proc Natl Acad Sci U S A, 110, 14314–14319.

61. Liu, X. and Roy, R. (2001) Mutation at active site lysine 212 to arginine uncouples the glycosylase activity from the lyase activity of human endonuclease III. Biochemistry, 40, 13617–13622.

62. Saito, Y., Uraki, F., Nakajima, S., Asaeda, A., Ono, K., Kubo, K. and Yamamoto, K. (1997) Characterization of endonuclease III (nth) and endonuclease VIII (nei) mutants of Escherichia coli K-12. J Bacteriol, 179, 3783–3785.

63. Golan, G., Zharkov, D.O., Feinberg, H., Fernandes, A.S., Zaika, E.I., Kycia, J.H., Grollman, A.P. and Shoham, G. (2005) Structure of the uncomplexed DNA repair enzyme endonuclease VIII indicates significant interdomain flexibility. Nucleic Acids Res, 33, 5006–5016.

64. Madeira, F., Park, Y.M., Lee, J., Buso, N., Gur, T., Madhusoodanan, N., Basutkar, P., Tivey, A.R.N., Potter, S.C., Finn, R.D. et al. (2019) The EMBL-EBI search and sequence analysis tools APIs in 2019. Nucleic Acids Res, 47, W636–W641.

65. Landau, M., Mayrose, I., Rosenberg, Y., Glaser, F., Martz, E., Pupko, T. and Ben-Tal, N. (2005) ConSurf 2005: the projection of evolutionary conservation scores of residues on protein structures. Nucleic Acids Res, 33, W299–302.

66. McCoy, A.J., Grosse-Kunstleve, R.W., Adams, P.D., Winn, M.D., Storoni, L.C. and Read, R.J. (2007) Phaser crystallographic software. J Appl Crystallogr, 40, 658–674.

67. Emsley, P., Lohkamp, B., Scott, W.G. and Cowtan, K. (2010) Features and development of Coot. Acta Crystallogr D Biol Crystallogr, 66, 486–501.

68. Zharkov, D.O., Golan, G., Gilboa, R., Fernandes, A.S., Gerchman, S.E., Kycia, J.H., Rieger, R.A., Grollman, A.P. and Shoham, G. (2002) Structural analysis of an Escherichia coli endonuclease VIII covalent reaction intermediate. EMBO J, 21, 789–800.

69. Eckenroth, B.E., Cao, V.B., Averill, A.M., Dragon, J.A. and Doublie, S. (2021) Unique Structural Features of Mammalian NEIL2 DNA Glycosylase Prime Its Activity for Diverse DNA Substrates and Environments. Structure, 29, 29–42 e24.

70. Parikh, S.S., Mol, C.D., Slupphaug, G., Bharati, S., Krokan, H.E. and Tainer, J.A. (1998) Base excision repair initiation revealed by crystal structures and binding kinetics of human uracil-DNA glycosylase with DNA. EMBO J, 17, 5214–5226.

71. Armougom, F., Moretti, S., Poirot, O., Audic, S., Dumas, P., Schaeli, B., Keduas, V. and Notredame, C. (2006) Expresso: automatic incorporation of structural information in multiple sequence alignments using 3D-Coffee. Nucleic Acids Res, 34, W604–608.

72. Kuznetsov, N.A., Koval, V.V., Nevinsky, G.A., Douglas, K.T., Zharkov, D.O. and Fedorova, O.S. (2007) Kinetic conformational analysis of human 8-oxoguanine-DNA glycosylase. J Biol Chem, 282, 1029–1038.

73. Kladova, O.A., Grin, I.R., Fedorova, O.S., Kuznetsov, N.A. and Zharkov, D.O. (2019) Conformational Dynamics of Damage Processing by Human DNA Glycosylase NEIL1. J Mol Biol, 431, 1098–1112.

74. Kuznetsov, N.A., Zharkov, D.O., Koval, V.V., Buckle, M. and Fedorova, O.S. (2009) Reversible chemical step and rate-limiting enzyme regeneration in the reaction catalyzed by formamidopyrimidine-DNA glycosylase. Biochemistry, 48, 11335–11343.

75. Kuznetsov, N.A., Kladova, O.A., Kuznetsova, A.A., Ishchenko, A.A., Saparbaev, M.K., Zharkov, D.O. and Fedorova, O.S. (2015) Conformational Dynamics of DNA Repair by Escherichia coli Endonuclease III. J Biol Chem, 290, 14338–14349.

76. Porter, C.M. and Miller, B.G. (2012) Cooperativity in monomeric enzymes with single ligand-binding sites. Bioorg Chem, 43, 44–50.

77. Kamata, K., Mitsuya, M., Nishimura, T., Eiki, J. and Nagata, Y. (2004) Structural basis for allosteric regulation of the monomeric allosteric enzyme human glucokinase. Structure, 12, 429–438.

78. Storer, A.C. and Cornish-Bowden, A. (1976) Kinetics of rat liver glucokinase. Co-operative interactions with glucose at physiologically significant concentrations. Biochem J, 159, 7–14.

79. Odell, I.D., Newick, K., Heintz, N.H., Wallace, S.S. and Pederson, D.S. (2010) Non-specific DNA binding interferes with the efficient excision of oxidative lesions from chromatin by the human DNA glycosylase, NEIL1. DNA Repair (Amst), 9, 134–143.

80. Robert, X. and Gouet, P. (2014) Deciphering key features in protein structures with the new ENDscript server. Nucleic Acids Res, 42, W320–324.

81. Landova, B. and Silhan, J. (2020) Conformational changes of DNA repair glycosylase MutM triggered by DNA binding. FEBS Lett, 594, 3032–3044.

82. Baba, D., Maita, N., Jee, J.G., Uchimura, Y., Saitoh, H., Sugasawa, K., Hanaoka, F., Tochio, H., Hiroaki, H. and Shirakawa, M. (2005) Crystal structure of thymine DNA glycosylase conjugated to SUMO-1. Nature, 435, 979–982.

83. Pidugu, L.S., Flowers, J.W., Coey, C.T., Pozharski, E., Greenberg, M.M. and Drohat, A.C. (2016) Structural Basis for Excision of 5-Formylcytosine by Thymine DNA Glycosylase. Biochemistry, 55, 6205–6208.

84. Zhu, C., Lu, L., Zhang, J., Yue, Z., Song, J., Zong, S., Liu, M., Stovicek, O., Gao, Y.Q. and Yi, C. (2016) Tautomerization-dependent recognition and excision of oxidation damage in base-excision DNA repair. Proc Natl Acad Sci U S A, 113, 7792–7797.

85. Doublie, S., Bandaru, V., Bond, J.P. and Wallace, S.S. (2004) The crystal structure of human endonuclease VIII-like 1 (NEIL1) reveals a zincless finger motif required for glycosylase activity. Proc Natl Acad Sci U S A, 101, 10284–10289.

86. Imamura, K., Wallace, S.S. and Doublie, S. (2009) Structural characterization of a viral NEIL1 ortholog unliganded and bound to abasic site-containing DNA. J Biol Chem, 284, 26174–26183.

87. Duclos, S., Aller, P., Jaruga, P., Dizdaroglu, M., Wallace, S.S. and Doublie, S. (2012) Structural and biochemical studies of a plant formamidopyrimidine-DNA glycosylase reveal why eukaryotic Fpg glycosylases do not excise 8-oxoguanine. DNA Repair (Amst), 11, 714–725.

88. Bruner, S.D., Norman, D.P. and Verdine, G.L. (2000) Structural basis for recognition and repair of the endogenous mutagen 8-oxoguanine in DNA. Nature, 403, 859–866.

89. Bjoras, M., Seeberg, E., Luna, L., Pearl, L.H. and Barrett, T.E. (2002) Reciprocal “flipping” underlies substrate recognition and catalytic activation by the human 8-oxo-guanine DNA glycosylase. J Mol Biol, 317, 171–177.

90. Faucher, F., Wallace, S.S. and Doublie, S. (2010) The C-terminal lysine of Ogg2 DNA glycosylases is a major molecular determinant for guanine/8-oxoguanine distinction. J Mol Biol, 397, 46–56.

91. Faucher, F., Duclos, S., Bandaru, V., Wallace, S.S. and Doublie, S. (2009) Crystal structures of two archaeal 8-oxoguanine DNA glycosylases provide structural insight into guanine/8-oxoguanine distinction. Structure, 17, 703–712.

92. Faucher, F., Robey-Bond, S.M., Wallace, S.S. and Doublie, S. (2009) Structural characterization of Clostridium acetobutylicum 8-oxoguanine DNA glycosylase in its apo form and in complex with 8-oxodeoxyguanosine. J Mol Biol, 387, 669–679.

93. Faucher, F., Wallace, S.S. and Doublie, S. (2009) Structural basis for the lack of opposite base specificity of Clostridium acetobutylicum 8-oxoguanine DNA glycosylase. DNA Repair (Amst), 8, 1283–1289.

